# Alternative splicing expands the functional portfolio of a plant virus to control the viral cycle

**DOI:** 10.64898/2026.02.19.706752

**Authors:** Delphine M. Pott, Laura Medina-Puche, Chaonan Shi, Jens Mülders, Hua Wei, Deborah Lapczinsky, Zara Yagci, Remila Ramasamy, Ying Li, Kyoka Kuroiwa, Björn Krenz, Linda Hanley-Bowdoin, Rosa Lozano-Durán

## Abstract

Viruses frequently have limited coding space, yet efficiently manipulate complex hosts by deploying intricate, often poorly understood strategies. Geminiviruses, devastating plant pathogens with small single-stranded (ss) DNA genomes, replicate in the nucleus aided by the viral replication-associated protein (Rep), the most conserved protein within this family and related ssDNA viruses. Rep initiates viral DNA replication and represses its own promoter to regulate the infection cycle. The molecular mechanisms enabling this dual functionality are so far unknown. Here, we show that the geminivirus tomato yellow leaf curl virus (TYLCV) exploits the host spliceosome to produce novel Rep splice variants. Splicing generates Rep isoforms lacking the central oligomerization domain that cannot initiate replication but strongly repress the Rep promoter. Conversely, Rep mutants deficient in splicing promote replication but fail to repress transcription. Reduced Rep splicing interferes with the viral gene expression hierarchy and decreases infectivity. Our findings therefore reveal a previously unrecognized viral strategy in which alternative splicing produces functionally specialized viral protein isoforms, providing a mechanistic explanation for the dual role of Rep in viral replication and gene regulation. Splicing events with potentially similar functional consequences in related viruses suggest that this strategy may have convergently evolved across diverse viral lineages infecting different domains of life.

## MAIN TEXT

Viruses are obligate intracellular parasites that frequently have limited coding space, but they are able to efficiently manipulate the much more complex organisms that they infect and facilitate their own multiplication and spread. The plant-infecting begomoviruses (family *Geminiviridae*) have circular single-stranded (ss) DNA genomes of 2.8-5.6 kb in size traditionally believed to encode only 6-8 proteins, yet they cause devastating diseases that can wipe out entire crop fields. To maximize their coding capacity, begomoviruses employ bidirectional transcription of overlapping genes (Schlub & Holmes, 2020; Chakraborty *et al*, 2022; Rojas *et al*, 2005). Even though they are transcribed in the nuclei of infected plant cells and have access to the splicing machinery, begomoviral transcripts were thought not to undergo splicing. However, recent results demonstrate that splicing is prevalent in the begomovirus tomato yellow leaf curl virus (TYLCV), and that it contributes to infection (Pott *et al*, 2025).

Strikingly, four of the eight splicing events identified in TYLCV transcripts affect the viral replication-associated (Rep) protein (Figure 1A; Pott *et al* (2025)). Rep is the largest TYLCV-encoded protein and the only one essential for replication of viral DNA. Rep binds the viral genome at a specific sequence located in the intergenic region through its N-terminal DNA-binding domain (Figure 1B), which results in two functional outcomes: on the one hand, Rep binding initiates replication by recruiting the host DNA replication machinery and nicking the virion DNA strand at the origin of replication; on the other hand, Rep also represses the activity of the Rep promoter to regulate progression through the infection cycle by enabling the transition from active viral replication to late functions, including movement and encapsidation (Rizvi *et al*, 2015; Ruhel & Chakraborty, 2019). It has been proposed that these two roles of Rep rely on independent processes (Eagle *et al*, 1994; Orozco *et al*, 1997), but how one viral protein can perform these two distinct roles through binding to the same sequence in the viral genome is poorly understood. Oligomerization of Rep through its central oligomerization domain (Figure 1A) seems to be specifically required for replication (Orozco *et al*, 2000).

**Figure 1.**
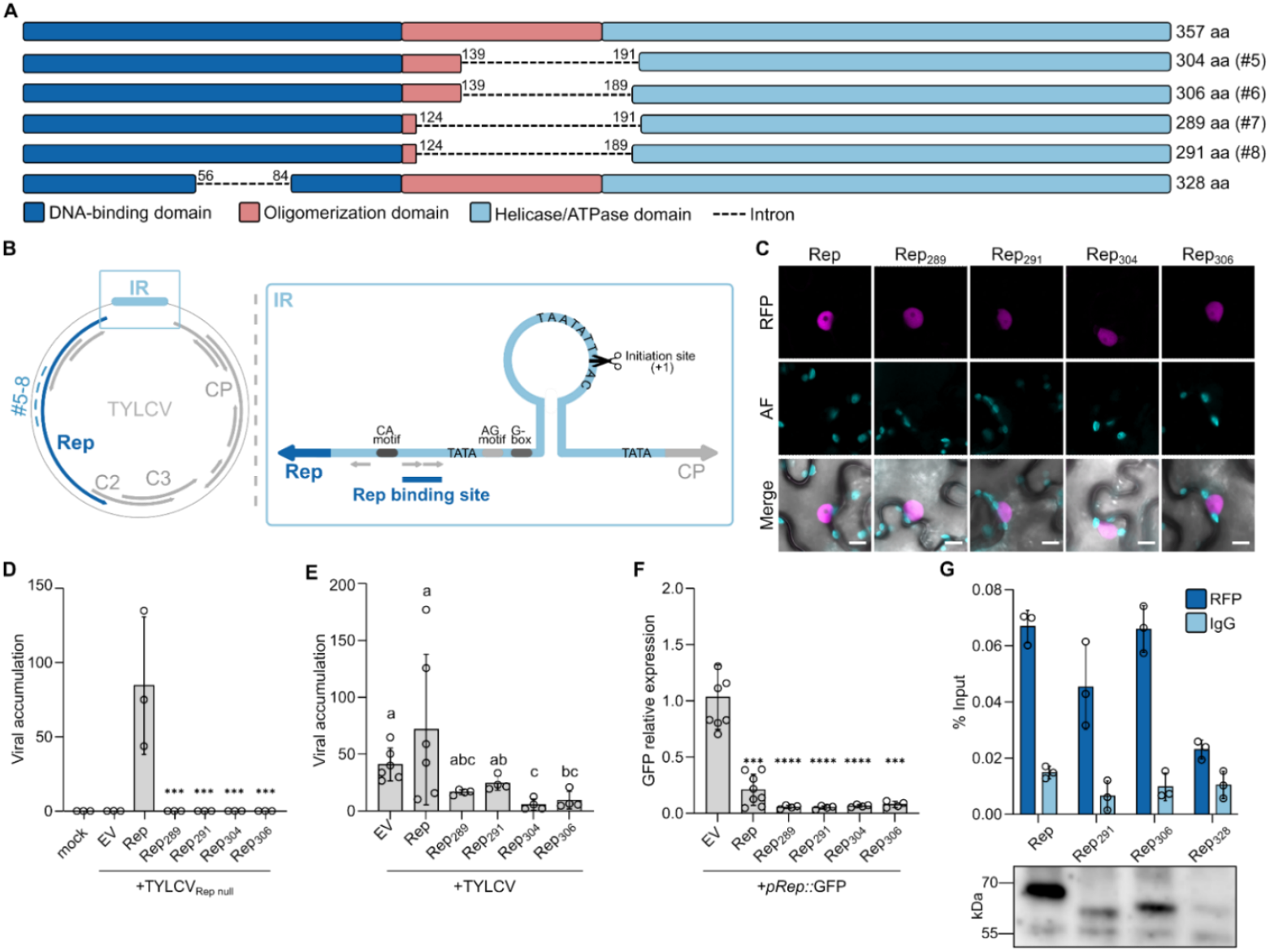
Rep splicing-derived isoforms lacking the oligomerization domain repress the Rep promoter. (A)Schematic representation of Rep and Rep splicing-derived isoforms. Four isoforms generated through alternative splicing lack the central oligomerization domain, while maintaining an intact C-terminal helicase domain, and yield proteins of 289, 291, 304, and 306 amino acids (aa), respectively. A fifth spliced isoform lacking part of the N-terminal DNA-binding domain, yielding a protein of 328 amino acids, with intact oligomerization and helicase/ATPase domains, was identified by Sanger sequencing. Splicing events identified in Pott *et al* (2025) are shown in parentheses (#5-8). (B)Schematic representation of the TYLCV gene organization. The positions of the Rep, C2, C3 and CP ORFs are indicated, as is the position of splicing events #5-8 (dashed line). A detailed zoom-in of the intergenic region (IR) is shown, with the stem-loop structure (including an invariant nonanucleotide sequence that contains the replication initiation site) and the Rep and CP promoter regions. Within the Rep promoter region, the Rep binding site is indicated as well as a series of motifs important for transcription and replication regulation (CA-, AG-motifs, G-box and TATA sequence). Grey arrows: iterated elements (iterons) that correspond to the Rep binding site. (C)Subcellular localization of Rep and Rep splicing-derived isoforms fused to C-terminal (Cter) RFP transiently expressed in *Nicotiana benthamiana*. Scale bar = 10 µm. (D)Rep splicing-derived isoforms do not complement a TYLCV Rep null mutant (Rep_null_) in local infection assays in *N. benthamiana*. Viral accumulation is measured by qPCR at 3 days post-infiltration (dpi). Error bars represent standard deviation (SD). The experiment was repeated twice with similar results (n=3-4). Results from one experiment are shown. A Dunnett post hoc test was used to compare splicing-derived isoforms to the full-length Rep with asterisks indicating significant differences. (E)Rep splicing-derived isoforms significantly decrease TYLCV accumulation in local infection assays in *N. benthamiana*. TYLCV infectious clone and the Rep different isoforms (expressed under the control of the 35S promoter) were co-infiltrated at the same time. Viral accumulation was measured by qPCR at 2 dpi. Error bars represent SD. The experiment was repeated three times with similar results (n=4-6). Results from one experiment are shown. Different letters above the bars indicate statistically significant differences at *p* < 0.05, based on Tukey’s post hoc test. (F)Rep splicing-derived isoforms strongly decrease GFP expression driven by Rep promoter (*pRep::*GFP). The Rep different isoforms (expressed under the control of the 35S promoter) and *pRep::*GFP were co-infiltrated at the same time. GFP expression is measured by qRT-PCR at 2 dpi. Error bars represent SD. The experiment was repeated three times with similar results (n=4-8). Results from one experiment are shown. Asterisks indicate statistically significant differences from EV samples as determined by Tukey’s post hoc test. (G)Rep splicing-derived isoforms (Rep_306_ and Rep_291_) bind to the Rep promoter in the TYLCV genome by ChIP-qPCR. The Rep different isoforms (expressed under the control of the 35S promoter) and *pRep::*Rep were co-infiltrated at the same time, and samples were harvested at 2 dpi and cross-linked with 1% formaldehyde. After nuclei extraction, anti-RFP antibody was used to immunoprecipitate the different RFP-tagged Rep isoforms, while mouse IgG antibody was included as a negative control. Rep_328_, which lacks part of the DNA-binding domain, is used as a negative control. Western blot of the input is shown. Error bars represent SD of three technical replicates. The experiment was repeated three times with similar results. Results from one experiment are shown.

Here, we show that RNA splicing of the Rep transcript can generate functionally specialized isoforms of the Rep protein. The full-length version of Rep promotes viral DNA replication but lacks transcriptional suppression activity, while splicing-derived variants lacking the central oligomerization domain are unable to mediate replication but strongly repress the Rep promoter. These results resolve the long-standing question of how the Rep protein coordinates these two functions and illustrate how splicing expands the functional portfolio of the viral proteome. Interestingly, independently evolved splicing events leading to removal of the central oligomerization domain of Rep proteins seem to be prevalent in phylogenetically related viruses infecting different domains of life, suggesting that the strategy followed by TYLCV to enable functional specialization of Rep might have convergently arisen across diverse hosts and viral lineages.

### Splicing-derived Rep isoforms lacking the oligomerization domain repress the Rep promoter

Five splicing events affecting the Rep transcript from TYLCV have been identified from RNA-seq data from locally infected *Nicotiana benthamiana* leaves (Wang *et al*, 2022; Pott *et al*, 2025). Four of them, resulting in removal of the central oligomerization domain of the Rep protein, were found using an automated pipeline (events #5-#8 in Pott *et al* (2025)), while a fifth one, producing a Rep isoform in which part of the DNA binding domain is missing, was identified by manual inspection of the reads and subsequently confirmed by Sanger sequencing (Figure 1A; Figure S1A). The Rep isoforms produced were named according to the size of the resulting protein (in amino acids) (Figure 1A).

Given that four of these splicing events remove the central oligomerization domain-coding sequence almost entirely with minimal differences, we decided to functionally characterize the resulting Rep isoforms (Rep_304_, Rep_306_, Rep_289_, and Rep_291_ corresponding to splicing events #5-8 in Pott *et al* (2025)). Prediction of the structure of these isoforms using AlphaFold3 (Abramson *et al*, 2024) suggests that they share high structural similarity. Specifically, the N-terminal part of the protein, which corresponds to the DNA binding domain, is unchanged compared to that in the original (full-length) Rep protein (see Table S1 for root mean square deviation values), while the C-terminal helicase domain is rotated due to loss of the central domain (Figure S1B; Table S1). All four Rep isoforms lacking the oligomerization domain localize to the nucleus, analogous to the full-length Rep, as shown by RFP or GFP fusions (Figure 1C; Figure S1C). However, none of them could mediate viral DNA replication in *N. benthamiana*, either in complementation assays with a TYLCV Rep null mutant (Figure 1D) or in a viral replicon-based system (*N. benthamiana* 2IR-GFP plants; Maio *et al*, 2019; Figure S1D). This lack of viral DNA replication-promoting activity is consistent with the inability of the new Rep isoforms to strongly oligomerize *in vivo*, with only a weak association detected in co-immunoprecipitation assays (Figure S1E, shown for Rep_291_ and Rep_306_ as representative of the four splicing-derived isoforms). Interestingly, self-interactions can be observed in yeast (Figure S1F), suggesting that a new interface for homotypic interactions might arise in the absence of the central oligomerization domain. These Rep isoforms seem to exert a dominant negative effect on viral accumulation in local infections (Figure 1E), which prompted us to test their capacity to repress the Rep promoter. Strikingly, we found that all four isoforms lacking the oligomerization domain can repress the Rep promoter, and this repression is stronger than that observed when Rep is expressed from its wild-type gene (Figure 1F). As expected, expression of these Rep isoforms has a negative impact on the production of viral replicons when the replication-competent full-length Rep is co-expressed from its own promoter, but not when it is expressed using a 35S promoter, indicating an effect at the level of Rep transcription (Figure S1G). Chromatin immunoprecipitation (ChIP) demonstrated that the Rep isoforms lacking the oligomerization domain (represented by Rep_291_ and Rep_306_) retain the ability to bind the Rep promoter (Figure 1G).

### TYLCV requires Rep isoforms lacking the oligomerization domain for full infectivity

To assess the contribution of the splicing-generated Rep isoforms lacking the oligomerization domain to infection, we generated mutant versions of Rep unable to produce these isoforms in the isolated gene or in the context of the viral genome. For this, the acceptor splice sites for splicing events #5-8 (producing Rep_304_, Rep_306_, Rep_289_, and Rep_291_) by themselves or with their respective donor sites were changed without affecting the full-length Rep protein sequence (Rep splicing mutant 1/#5-8_Δas_ (Pott *et al*, 2025) and mutant 2/#5-8_Δd,as_, respectively; Figure 2A; Figure S2A). Unexpectedly, mutation of the shared acceptor splice site for the splicing events #6 and #8 led to the generation of a novel, alternative splice site (Figure S2A-D) that resulted in the production of an artificial isoform of Rep similar to Rep_291_ and Rep_306_ but with a 1-bp shift, producing a truncated protein of 141 amino acids (Figure S2C-D). This was not the case when both donor and acceptor sites were mutated (Figure S2C). In line with this observation, Rep protein accumulation was decreased in splicing mutant 1 (Rep_/#5-8Δas_) compared to mutant 2 (Rep_#5-8Δd,as_) or wild-type Rep when transiently expressed in *N. benthamiana* fused to RFP (Figure S2E).

**Figure 2.**
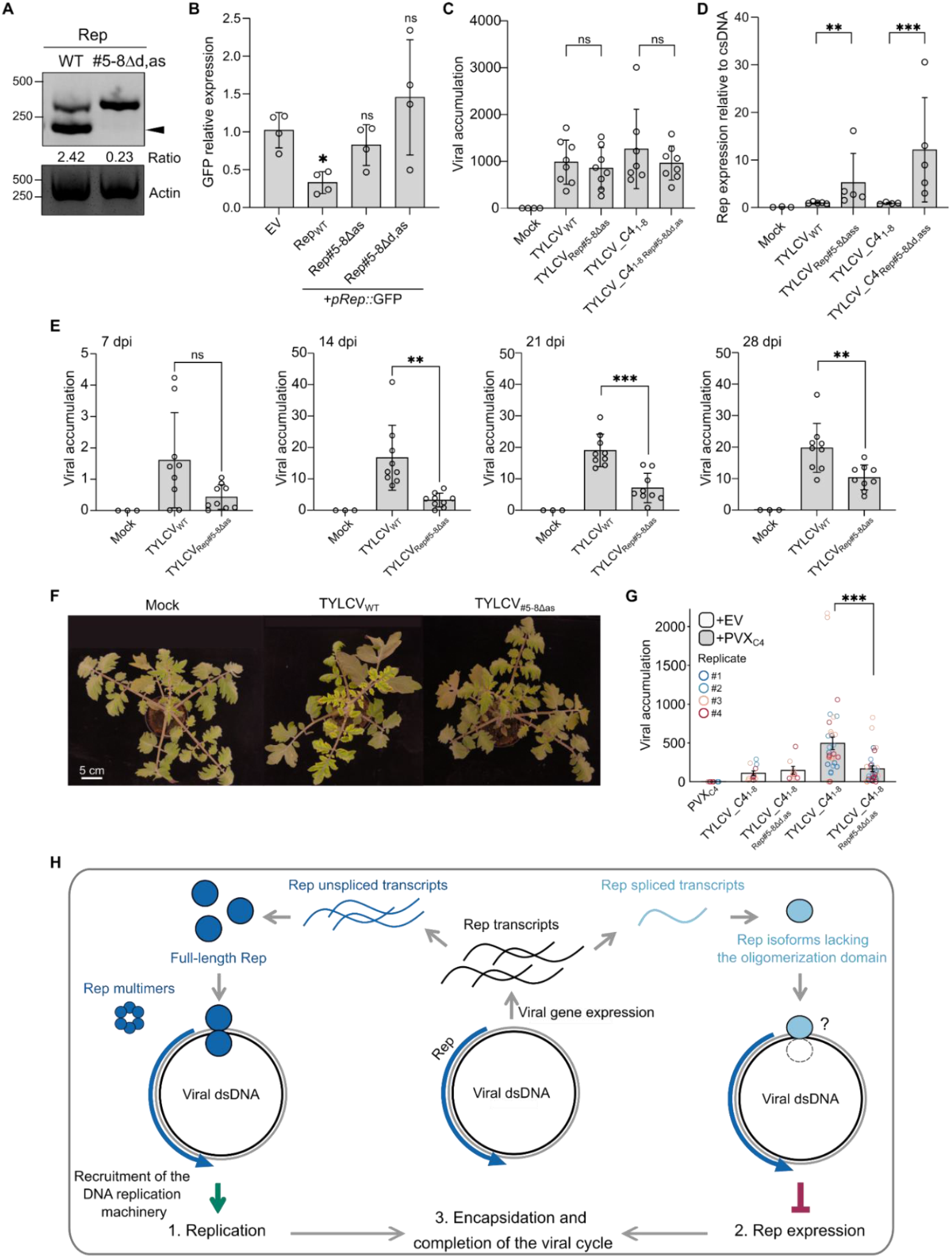
Splicing-derived Rep isoforms lacking the oligomerization domain are required for full infectivity. (A)Agarose gel showing cDNA from Rep_WT_ and Rep_#5-8Δd,as_-expressing samples. Used primers allow the amplification of the full-length Rep and Rep_304_ (Pott *et al*, 2025). Spliced-to-unspliced transcript ratio measured with ImageJ is shown under the gel image. The arrowhead indicates the spliced transcript in the agarose gel. (B)Rep mutants impaired in the four splicing events affecting the oligomerization domain (Rep_#5-8Δas_ and Rep_#5-8Δd,as_) lose the capacity to repress the Rep promoter, as shown by *GFP* expression driven by Rep promoter (*pRep::*GFP) and co-expressed either with an empty vector (EV) as a negative control, Rep_WT_, Rep_#5-8Δas_ or Rep_#5-8Δd,as_. GFP expression is measured by qRT-PCR at 2 dpi. Error bars represent SD. The experiment was repeated three times with similar results (n=4). Results from one experiment are shown. Asterisks indicate statistically significant differences from EV samples as determined by Dunnett T3 post hoc test; ns: non-significant. (C)TYLCV Rep mutants impaired in the four splicing events (TYLCV_Rep#5-8Δas_ and TYLCV_C4_1-8 Rep#5-8Δd,as_) affecting the oligomerization domain retain the capacity to replicate in local infection assays in *N. benthamiana*. Viral accumulation is measured by qPCR at 3 dpi. Error bars represent SD. The experiment was repeated three (for TYLCV_C4_1-8 Rep#5-8Δd,as_) and four (for TYLCV_Rep#5-8Δas_) times with similar results (n=6-8, see Figure S2H for Rep_#5-8Δas_ for the different replicates). Results from one experiment are shown. Statistical analysis is shown after Tukey post hoc test. (D)Rep expression is increased in TYLCV_Rep#5-8Δas_ and TYLCV_C4_1-8 Rep#5-8Δd,as_. Rep expression is normalized to the complementary strand viral DNA (csDNA) and is measured in infected samples from (C). Error bars represent SD. The experiment was repeated three times with similar results (n=6). Results from one experiment are shown. Asterisks indicate statistically significant differences as determined by Tukey’s post hoc test. (E)TYLCV_Rep#5-8Δas_ shows decreased viral accumulation in systemic infection assays in tomato (7-28 dpi). Error bars represent SD. The experiment was repeated four times with similar results (n=6-9). Results from one experiment are shown. Asterisks indicate statistically significant differences between TYLCV_WT_ and TYLCV_Rep#5-8Δas_ as determined by Tukey’s post hoc test. Cautionary note: the y-axis scale differs for 7 dpi. (F)Symptoms of the plants from (E) at 26 dpi. Scale bar = 5 cm. (G)TYLCV_C4_1-8 Rep#5-8Δd,as_ shows decreased viral accumulation in systemic infection assays (14 dpi) in *N. benthamiana*, compared to TYLCV_C4_1-8_. The lack of C4 protein was complemented by expression of the corresponding viral gene from a PVX-based vector (PVX_C4_). Error bars represent SE. The experiment was repeated four times with similar results (n=7-9). A linear mixed-effects model with the genotypes as a fixed effect and the replicates as a random effect was used. Asterisks indicate statistically significant differences after pairwise comparisons among groups using estimated marginal means with Tukey-adjusted p-values. (H)Model showing splicing of Rep transcripts as a mechanism to produce functionally-specialized protein isoforms. Rep is the first gene to be expressed from a double-stranded (ds) DNA intermediate of the viral genome (viral dsDNA) in the infected plant nucleus. A small proportion of Rep transcripts is alternatively spliced, producing Rep isoforms lacking the central oligomerization domain (Pott *et al*, 2025). The main isoform, corresponding to the full-length Rep, self-interacts and binds to the viral genome in the intergenic region to recruit the host DNA replication machinery and initiate viral genome replication. Rep also forms high order multimers, a step required for its role in replication (Orozco *et al*, 2000). The splicing-derived Rep isoforms are also able to bind the same site in the intergenic region, where the Rep promoter is localized. This binding is not productive for replication, likely due to the inability of the isoforms to form replication-competent homotypic complexes, and results in the transcriptional repression of the Rep gene. The activity of these Rep isoforms, which “poison” the origins of replication they bind, favors the expression of viral late genes, hence potentially promoting the transition in the viral cycle from active replication to viral movement and encapsidation.

We reasoned that both the full-length version of the protein as well as its spliced isoforms are likely produced from the wild-type Rep gene, as previously decribed in the context of the infection (Pott *et al*, 2025). Two observations support this hypothesis: i) both Rep splicing mutants that cannot produce the Rep isoforms lacking the oligomerization domain are unable to suppress the Rep promoter (Figure 2B); and ii) the infectious clones carrying Rep splicing mutants are able to replicate in local viral infections in *N. benthamiana* (Figure 2C; Figure S2H), with the resulting Rep protein mediating replication of a viral replicon (Figure S2F-G). The release of Rep promoter repression was also observed when viral transcript levels were measured during local viral infection (Figure 2D). These results strongly suggest that splicing leads to functionally specialized Rep isoforms, with the full-length protein promoting viral DNA replication and the oligomerization domain-deficient isoforms repressing the Rep promoter.

A TYLCV infectious clone containing the Rep_#5-8Δas_ mutation displayed reduced infectivity in tomato plants (Figure 2E, F), as previously observed in *N. benthamiana* (Pott *et al*, 2025). Mutating the donor splice sites unavoidably affects the sequence of the viral C4 protein encoded by the overlapping C4 gene. To address any potential effect of the C4 mutation, we introduced Rep splicing mutant 2 (Rep#_5-8Δd,as_) in a C4 null mutant background (TYLCV_C4_1-8_, Rosas-Diaz *et al* (2018)), generating TYLCV_C4_1-8 Rep#5-8Δd,as_, and complemented the C4 mutation by expressing the corresponding gene from a potato virus X (PVX)-based vector (Icon Genetics GmbH). As previously described, TYLCV_C4_1-8_ could not infect tomato plants (Rosas-Diaz *et al*, 2018) and shows low infectivity in *N. benthamiana* (Figure 2G). Co-inoculation with a PVX clone containing the C4 sequence (PVX_C4_) led to a partial complementation of the TYLCV_C4_1-8_ mutant but did not affect accumulation of TYLCV_C4_1-8 Rep#5-8Δd, as_ (Figure 2G), demonstrating a specific penalty to the infection process attributable to the missing splicing events. Taken together, the results of these infection experiments indicate that splicing events removing the central oligomerization domain in Rep are required for full viral infectivity but do not impair viral replication.

Rep-mediated repression of its own promoter is hypothesized to be necessary for downregulating Rep expression at the appropriate time in the viral infection cycle, thereby permitting expression of late viral genes and the shift from active replication to genome movement and encapsidation (Shung & Sunter, 2007; Borah *et al*, 2016). To evaluate the effect of splicing-derived Rep isoforms on the expression of viral genes, we first measured the transcript accumulation of the C2 and C3 genes, located downstream of Rep in the complementary strand of the viral genome, from a partial genome comprising a fragment from the intergenic region (containing the Rep promoter) to the end of the C3 gene (Figure 1B; Figure S2I). This partial viral genome contains mutations in the Rep gene that render it unable to produce the corresponding protein (as in Figure 1D, Rep null mutant). Transcript levels were measured after *in trans* expression of Rep spliced isoforms Rep_291_ and Rep_306_, or full-length Rep from its wild-type gene or the splicing mutant 2 (Rep_/#5-8Δd,as_). Surprisingly, the presence of full-length Rep leads to an increase in the Rep and C2/C3 transcripts. This effect is consistently more pronounced when Rep is produced from the splicing mutant. In contrast, expression of early and intermediate viral genes (Rep, C2, and C3) was repressed in the presence of the Rep_291_ and Rep_306_ isoforms (Figure S2I). Analysis of the expression of CP, a late viral gene encoding the capsid protein, in infected plants from Figure 2G demonstrated that the absence of Rep isoforms lacking the oligomerization domain leads to a reduction in the CP transcript (Figure S2J). This decreased expression of the late gene may account for the reduction in infection observed for the tested splicing mutants (Figure 2E-G), and supports the notion that autoregulatory Rep expression is required to establish the hierarchy of viral gene expression, in accordance with previous models (Shung & Sunter, 2007; Borah *et al*, 2016). Taken together, our results illustrate how alternative splicing generates a set of novel isoforms of the Rep protein that repress the expression of its own gene and orchestrate the hierarchy of viral gene expression during infection (Figure 2H).

### Splicing-enabled removal of the central oligomerization domain in Rep occurs in ssDNA viruses across kingdoms

Given that the dual roles of Rep have been described in geminiviral species other than TYLCV (Eagle *et al*, 1994; Fontes *et al*, 1992; Haley *et al*, 1992; Borah *et al*, 2016), we first sought to determine whether splicing of the Rep-coding transcript is a common feature within the genus *Begomovirus* in the family *Geminiviridae*. Prediction tools identified high-confidence splicing events removing the central oligomerization domain-coding sequence in all tested begomoviruses across different lineages, i.e. begomoviruses from the Africa, Asia, Europe and Oceania (AAEO) region, begomoviruses form the Americas, and sweet potato-infecting begomoviruses (Figure 3A-B; Table S2). Conservation of putative splice sites (either donor or acceptor) is widespread among begomoviruses, although the same introns as in TYLCV could be identified only in some AAEO-region species. The predicted splicing events would result in one of two main outcomes at the protein level: 1) the removal of the central domain while maintaining the N- and C-terminal domains, as shown for TYLCV (e.g. squash leaf curl virus (SLCuV), sweet potato leaf curl Henan virus (SPLCHnV), or tomato leaf curl Joydebpur virus (ToLCJV)); or 2) a frameshift leading to a premature stop codon, which would either maintain the complete DNA binding domain (e.g. East African cassava mosaic virus (EACMV), tomato leaf deformation virus (ToLDeV) or Jatropha mosaic virus (JMV)) or truncate it after the first 85-90 amino acids (e.g. tomato yellow leaf curl China virus (TYLCCNV), tomato leaf curl purple vein virus (ToLCPVV) or chilli leaf curl virus (ChiLCV)). We experimentally confirmed the splicing event detected in the Rep transcript from the bipartite begomovirus EACMV, representing the second case (Figure 3C, D; Figure S3). Interestingly, these results suggest that splicing events leading to Rep isoforms belonging to one of two classes (TYLCV-like and EACMV-like) have likely independently evolved several times within the begomovirus clade.

**Figure 3.**
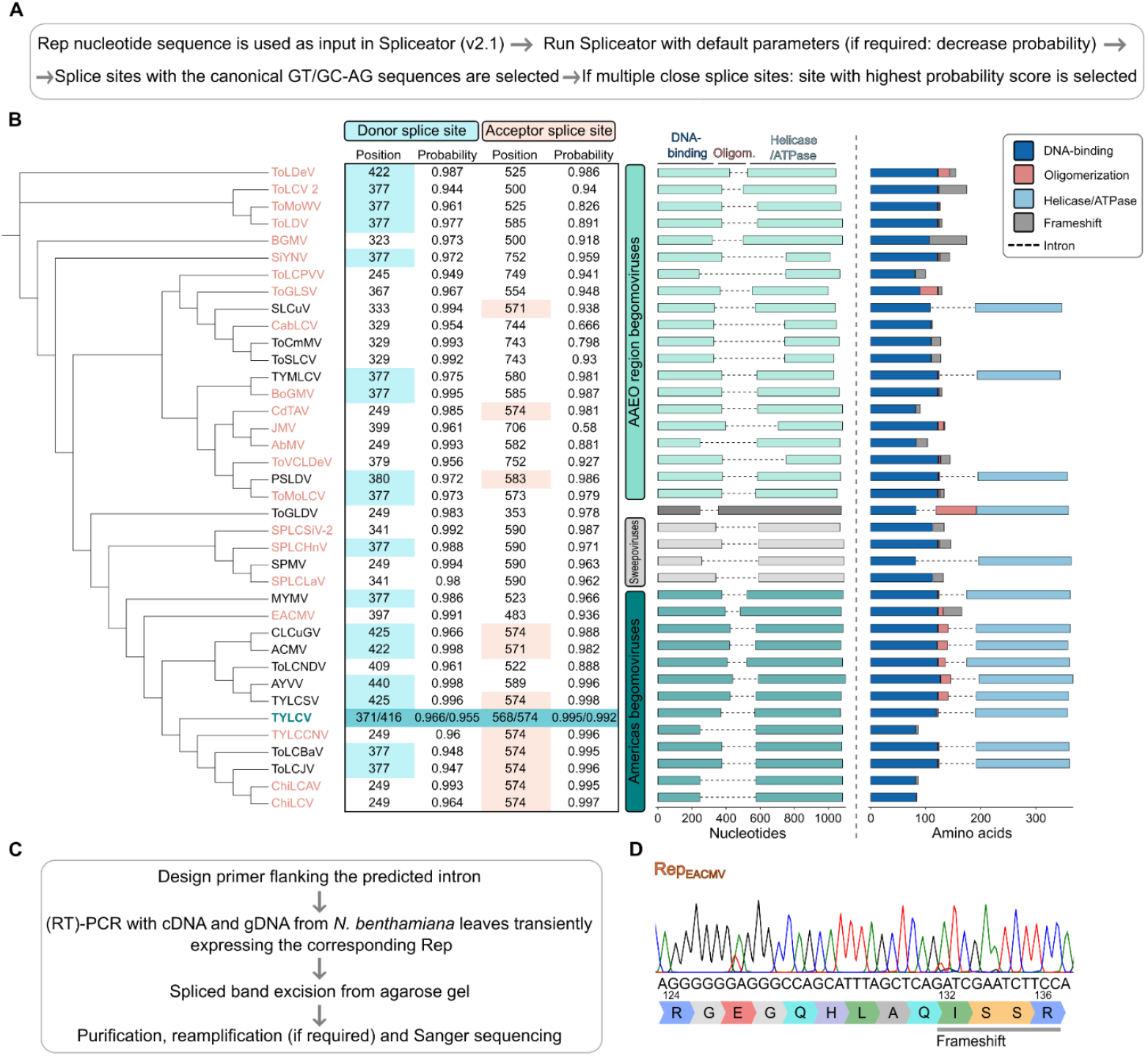
Splicing of Rep transcript occurs in begomoviruses from different lineages. (A)Pipeline for the prediction of splice sites using Spliceator predictor tool (Scalzitti *et al*, 2021) (B)Phylogeny (inferred from Rep amino acid sequence with MUSCLE (EMBL-EBI)) and splicing predictions of the central region of Rep transcripts in selected begomoviruses. Donor and acceptor splice site positions and prediction scores are shown. Splice sites conserved with those of TYLCV are highlighted in blue and orange, respectively. Schematics depict the predicted spliced transcripts and resulting protein isoforms. Rep domains were assigned by alignment with TYLCV Rep (Figure S5). Viruses’ names in orange indicate predicted truncated protein isoforms. Virus full names, phylogeny and accession numbers are listed in Table S2. AAEO: Africa, Asia, Europe, Oceania; Sweepoviruses: sweet potato-infecting begomoviruses; Oligom.: Oligomerization. (C)Pipeline for the experimental confirmation of splicing events detected in Rep transcripts. (D)Sanger sequencing for the detection of the spliced isoform of Rep transcript from EACMV.

Geminiviruses belong to the Circular Rep-encoding Single-Stranded DNA (CRESS) phylum, which is comprised of families that are defined by encoding a Rep protein and using the same mode of replication (Rep-enabled rolling circle) but can infect different domains of life (Figure 4A). Considering that the need to transition from a phase of active genome replication to encapsidation is likely a universal imperative of viral life cycles, we investigated whether similar splicing events affecting the Rep transcript exist in other CRESS viruses. Notably, two splicing events resulting in the removal of the central part of the Rep transcript have been experimentally identified in the circovirus porcine circovirus 2 (PCV2) (Figure 4B; Bratanich & Blanchetot (2002)), although they could not be predicted (Figure 4A). Splicing events that lead to the full or partial removal of the Rep oligomerization domain, analogous Rep splicing in TYLCV and other begomoviruses, can also be predicted in other geminiviral genera as well as in pea necrotic yellow dwarf virus (PNYDV), a member of the *Nanoviridae* family, the only other CRESS family infecting plants (Figure 4A-B; Table S2). The splicing event predicted for the nanovirus PNYDV was also experimentally confirmed (Figure 4C; Figure S4).

**Figure 4.**
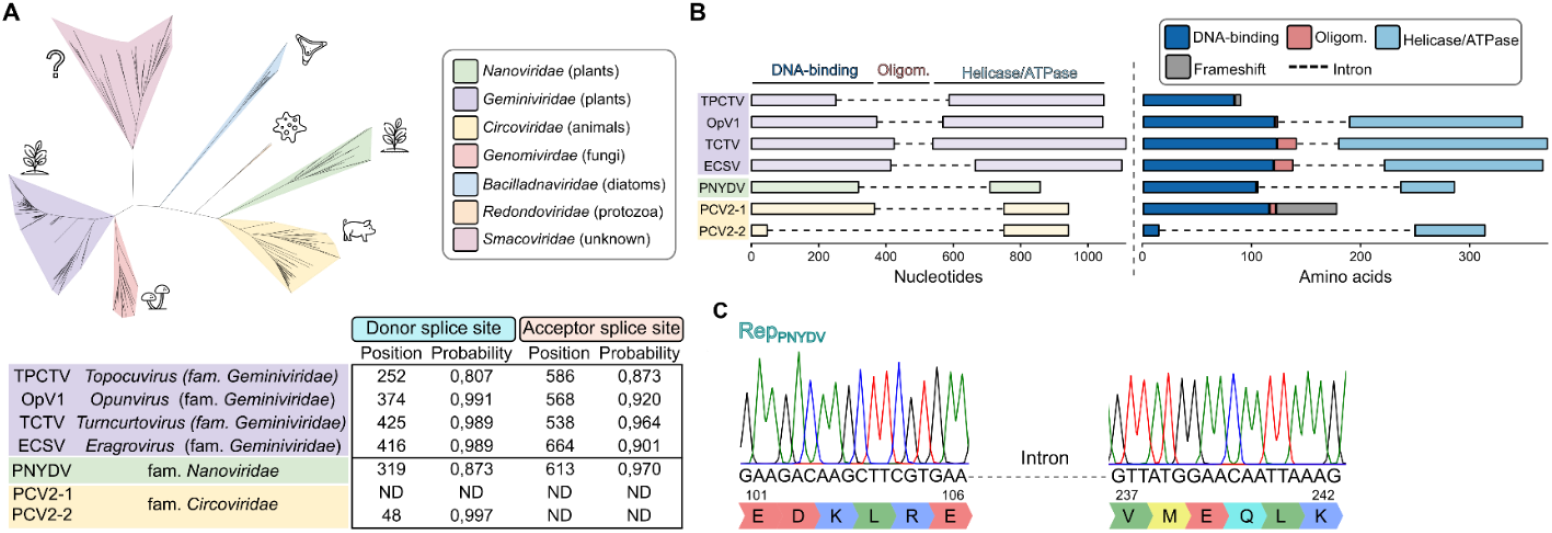
Splicing of Rep leading to the removal of the oligomerization domain coding sequence also occurs in other ssDNA viruses. (A)Splicing prediction of the central region of Rep transcripts in other genera of geminiviruses (*Topocuvirus, Opunvirus, Turncurtovirus* and *Eragrovirus*) and other CRESS viruses (pea necrotic yellow dwarf virus; PNYDV, fam. *Nanoviridae*). Two splicing events in Rep from porcine circovirus 2 (PCV2, fam. *Circoviridae*) were previously reported (Bratanich & Blanchetot, 2002). A phylogenetic tree of Rep proteins from representative eukaryote-infecting circular Rep-encoding single-stranded (CRESS) DNA viruses is shown. Amino acid sequences were obtained from Kazlauskas *et al* (2019) and supplemented with updated entries retrieved from the International Committee on Taxonomy of Viruses (ICTV) and the NCBI database. Multiple sequence alignment was performed in Jalview using MUSCLE, and the phylogenetic tree was constructed by the neighbor-joining method with 1,000 bootstraps, using phylogeny.fr (Dereeper *et al*, 2008) and edited with iTOL5 (Letunic & Bork, 2021). ND: not detected. (B)Schematic representation of the predicted spliced transcripts and resulting protein isoforms from A. Rep domains were assigned by alignment with TYLCV Rep for geminiviruses (Figure S6A), or were calculated based on Alphafold3 prediction (for PNYDV) or available structural data (for PCV2, Tarasova *et al* (2021)) of the corresponding Rep structure (Figure S6B). Virus full names and accession numbers are listed in Table S2. Oligom.: Oligomerization. (C)Sanger sequencing for the detection of the spliced isoform of Rep transcript from PNYDV.

These results suggest that splicing of the Rep transcript may be a general, convergently evolved strategy to diversify Rep functions and ensure the coordinated progression of the viral life cycle across CRESS viruses.

### Splicing of Rep as a mechanism to produce functionally specialized viral protein isoforms and facilitate the switch from replication to later phases of the infection cycle

Although prevalent splicing in TYLCV has been experimentally confirmed and the relevance of one of these events for the infection demonstrated (Pott *et al*, 2025), whether splicing of viral transcripts results in protein isoforms with distinct functions remained to be determined. Here, we show that alternative splicing of TYLCV Rep transcripts produces new versions of the protein that acquire the ability to repress the expression of the Rep-coding gene, demonstrating that splicing facilitates the functional expansion of the viral proteome (Figure 2H).

Spliced Rep transcripts are significantly less abundant than unspliced ones (Pott *et al*, 2025). However, the resulting protein isoforms devoid of the central oligomerization domain exert a dominant-negative effect over the replication activity of the full-length Rep (Figure 1E). Although these isoforms are likely produced at low levels, their properties make them highly efficient negative regulators of TYLCV replication. They bind the same sequence as full-length Rep but are unlikely to efficiently form the multimeric Rep complexes required for replication of the viral genome, based on the weak self-interaction detected in vivo (Figure S1E) (Orozco *et al*, 2000; Clérot & Bernardi, 2006; Tarasova *et al*, 2021; Boer *et al*, 2009). Each binding event may “poison” an origin of replication by preventing assembly of the functional Rep homotypic complexes. This nonproductive occupancy concomitantly represses transcription of the Rep gene, reducing the production of full-length Rep in the infected cell. As more Rep-binding sequences become blocked, fewer genomes remain capable of producing Rep, raising the relative impact of the splicing-derived isoforms and creating a self-reinforcing decline in replication competence. Over time, this feedback loop would allow even a rare protein variant to progressively lead to replication collapse. This Rep spliced variants-triggered replication collapse could be analogous to the effect observed when a truncated version of Rep that is capable of binding to the viral genome and repressing Rep gene expression is artificially expressed and confers resistance to the viral infection by inhibiting viral replication (Lucioli *et al*, 2003; Sardo *et al*, 2011). An RGG motif at the beginning of the oligomerization domain conserved in Rep from begomoviruses was proposed to be essential for this effect (Sardo *et al*, 2011). Notably, the spliced variants characterized in this work (Figure 1A) and most of the predicted ones (Figure 3B; Figure S5) contain and maintain the initial RG of this motif, supporting the involvement of these residues in transcriptional repression. The shutting down of viral replication and repression of early viral genes are prerequisites for the expression of late viral genes that are essential for movement and encapsidation of the virus. Therefore, the “origin poisoning” by the splicing-derived Rep isoforms is presumably also a requirement for time-controlled late gene expression and essential for full infectivity. This notion is supported by the positive effect of Rep isoforms on CP transcript accumulation (Figure S2J) as well as by the differential penalty of preventing production of the Rep splicing-derived isoforms on viral replication (Figure 2C; Figure S2F-H) and systemic infection (Figure 2E-G).

Enticingly, the strategy of generating Rep isoforms with dominant-negative effects on viral replication may have evolved independently across multiple CRESS viral families infecting hosts from different kingdoms of life, as suggested by the experimental or predicted identification of spliced Rep transcripts lacking the coding region for the central oligomerization domain (Figures 3-4). Such convergence underscores the potential efficacy of this regulatory design.

Reinforcing this idea, phylogenetically unrelated DNA viruses infecting animals, such as adenoviruses and polyomaviruses, appear to regulate their cycles via similar strategies. In adenoviruses, E1A is the first viral gene expressed and acts as a master transactivator of early viral gene expression, while also remodelling the cellular environment to become replication-permissive (Radko *et al*, 2014). Alternative splicing generates two principal mRNAs, 13S and 9S, that are predominant during the early and late phases of infection, respectively. The 13S isoform is a potent activator of early viral transcription and drives host cells into S phase, boosting the availability of the DNA replication machinery required at infection onset. The 9S isoform, a comparatively weaker transactivator, modulates transcription over time and facilitates the transition to late-phase viral gene expression.

A comparable splicing-based regulatory strategy also operates in polyomaviruses. Their early region produces several isoforms of the tumor antigens through alternative splicing, most prominently the large (LT) and small (ST) T antigens. LT is essential for viral DNA replication and, as its levels accumulate, it represses early gene transcription and promotes the expression of late genes, thereby coordinating the switch from genome amplification to virion assembly. ST, while multifunctional, modulates host signalling pathways that enhance early viral gene expression and support establishment of infection (Topalis *et al*, 2013).

Taken together, our results demonstrate that alternative splicing of viral transcripts constitutes a powerful regulatory layer in geminivirus biology, enabling small plant-infecting DNA viruses to diversify protein function and orchestrate a tightly controlled progression from replication to encapsidation. Future functional dissection of the splicing-derived viral protein isoforms will be fundamental for fully understanding – and ultimately disrupting – the geminivirus infection cycle.

## MATERIALS AND METHODS

### Plant material

*Solanum lycopersicum* cv. “Moneymaker”, *Nicotiana benthamiana* wild type and 2IR-GFP *N. benthamiana* transgenic lines (described in Maio *et al* (2019)) were grown in a controlled growth chamber under long-day conditions (16 h of light/8 h of dark) at 25 °C.

### Bacterial strains and growth conditions

*Escherichia coli* strain DH5α was used for general cloning and subcloning procedures; strain DB3.1 was used to amplify Gateway-compatible empty vectors. *Agrobacterium tumefaciens* strain GV3101 harboring the corresponding binary vectors was used for *in planta* expression.

### Plasmid construction

All primers and plasmids used for cloning are summarized in Tables S3 and S4, respectively. The TYLCV isolate used as a template is AJ489258 (NCBI:txid220938, as described in Wang *et al*, 2017; Rosas-Diaz *et al*, 2018). The TYLCV_C4_1-8_ and TYLCV_Rep#5-8Δas_ infectious clones are described in Rosas-Diaz *et al* (2018) and Pott *et al* (2025), respectively. The TYLCV Rep null mutant was synthesized, converting the first, fourteenth, fifty-third and sixty-third codons (encoding methionine, leucine and two glutamates, respectively) to stop codons (ATG → TAG, CTA → TAA, GAA → TAA and GAA → TAA). TYLCV_C4_1-8 Rep#5-8Δd,as_ mutant was generated by site-directed mutagenesis PCR, with the corresponding primers (Table S3). The DNA fragments carrying the mutations were then cloned into the pDONR/Zeo entry vector (Thermo Scientific) and were further recombined into pGWB501 destination vector through a Gateway LR reaction (Thermo Scientific).

Rep spliced variants (Rep_289_, Rep_291_, Rep_304_, Rep_306_ and Rep_328_) were generated by overlap PCR to remove the spliced fragments from Rep coding sequence and were cloned into pDONR/Zeo entry vector with or without a stop codon, to enable N- or C-terminal protein fusions, respectively. To generate the construct used for the analysis of Rep promoter activity, the TYLCV intergenic region in the complementary strand orientation was PCR-amplified and cloned into pDONR/Zeo vector. The sequence was then introduced into pGWB504 by a Gateway LR reaction to generate *pRep::*GFP. To generate *pRep::*Rep_null_-C2-C3, the TYLCV intergenic region in the complementary strand sense was amplified together with the corresponding Rep, C2 and C3 open reading frames and introduced into pGWB501 by a Gateway LR reaction. Vectors from the pGWB series are described in Nakagawa *et al* (2007a, 2007b).

For yeast two-hybrid (Y2H) assays, Rep_291_ and Rep_306_ were introduced into pGBKT7-GW and pGADT7-GW by Gateway cloning (Lu *et al*, 2010), while Rep full length was introduced into Golden Gate-compatible vectors, pGTT2t7AD and pGTT2t7BK, using Esp3I cut-ligation.

For PVX_C4_ complementation, the C4 sequence from TYLCV was cloned into pICH31160 (Icon Genetics) by Golden Gate cloning.

### *Agrobacterium*-mediated transient gene expression in *N. benthamiana*

For local infiltrations, *A. tumefaciens* cells carrying the constructs of interest were liquid-cultured in LB with the appropriate antibiotics at 28°C overnight. Bacterial cultures were centrifuged at 4,000 × *g* for 5 min and resuspended in infiltration buffer (10 mM MgCl_2_, 10 mM MES, pH 5.6, and 150 µM acetosyringone) to an OD_600_ = 0.1–0.5. After a 2-h incubation at room temperature in the dark, bacterial cultures were used to infiltrate the abaxial side of leaves of 3-to 4-week-old *N. benthamiana* wild-type or 2IR-GFP plants with a 1 mL needleless syringe. For experiments that required co-infiltration, the bacterial suspensions carrying different constructs were mixed at 1:1 ratio before infiltration. Plants were kept in the greenhouse for 24-48 h before further analysis.

### TYLCV infection

For TYLCV local infection assays, fully expanded young leaves of 4-week-old *N. benthamiana* plants were infiltrated with *A. tumefaciens* carrying TYLCV infectious clones (TYLCV wild type, TYLCV_Rep null_, TYLCV_C4_1-8_, TYLCV _Rep#5-8Δas_, or TYLCV_C4 1-8 #5-8Δd,as_). Samples were collected at 3 days post-inoculation (dpi) to detect viral accumulation and viral gene expression. For TYLCV systemic infection assays, the stems of 2-week-old tomato plants were syringe-inoculated with *A. tumefaciens* carrying TYLCV infectious clones (TYLCV wild type, TYLCV _Rep#5-8Δas_). The second youngest apical leaf was harvested at 7-28 dpi to detect viral accumulation. For complementation with PVX_C4_, cultures harboring TYLCV and PVX_C4_ infectious clones were mixed at a 1:1 ratio. The stem and the first two true leaves of 2-week old *N. benthamiana* plants were syringe-inoculated. The two youngest leaves were harvested at 14 dpi.

### Determination of viral accumulation and quantitative PCR (qPCR)

To determine viral accumulation, total DNA was extracted using the CTAB method (Murray & Thompson, 1980). The DNA from local infection assays was treated with FastDigest *Dpn*I (NEB) at 37°C for 30 min prior to further analysis. Quantitative PCR (qPCR) was performed with primers to amplify TYLCV Rep (Wang *et al*, 2017). As internal reference for DNA detection, the *25S ribosomal DNA interspacer* (*ITS*) was used (Mason *et al*, 2008). qPCR was performed in a BioRad CFX384 real-time system with PowerTrack SYBR Green Mastermix (Thermo Scientific), following the program: 2 min at 95°C, and 40 cycles consisting of 15 s at 95°C, 1 min at 60°C. For viral gene expression, complementary-strand (cs) DNA was amplified from 100 ng of total DNA with T4 DNA polymerase (ThermoFisher). After PCR purification, csDNA was measured by qPCR as described above and with specific primers (Table S3).

To determine the accumulation of GFP replicons in 2IR-GFP plants, total DNA was analyzed by qPCR with primers amplifying *GFP*, and *ITS* as a reference gene. All the primers used are described in Table S3.

### RNA extraction and Reverse transcription quantitative PCR (RT-qPCR)

RNA was extracted following the citrate-citric acid RNA isolation method (Oñate-Sánchez & Verdonk, 2021). RNA was treated with DNAse I (Thermo Scientific), and 500 ng of clean RNA was used for cDNA synthesis, with the PrimeScript RT MasterMix (Takara) following the manufacturer’s instructions. The qPCR reaction was performed with PowerTrack SYBR Green Mastermix, following the program: 2 min at 95°C, and 40 cycles consisting of 15 s at 95°C, 1 min at 60°C. *Elongation factor-1 alpha* (*NbEF1α*) (Segonzac *et al*, 2011) was used as reference gene. The primers used are described in Table S3.

### Visualization of subcellular localization with confocal microscope

For subcellular localization, *N. benthamiana* leaves expressing GFP-or RFP-fused proteins were imaged with a Confocal Laser Scanning Platform Leica TCS SP8 (Leica) or a LSM880 Airyscan confocal microscope (Zeiss). The preset settings for GFP (Ex: 488 nm, Em: 500–550 nm) or RFP (Ex: 554 nm, Em: 580-630 nm) were used.

### GFP fluorescence measurement

Leaf discs (5 mm diameter) were collected and placed adaxial side down in a black 96-well plate with 300 μL distilled water per well. Fluorescence (excitation 490 nm, emission 510– 570 nm) was measured using a Berthold Mithras LB940 plate reader.

### Protein extraction and co-immunoprecipitation (co-IP) assays

*A. tumefaciens* clones harboring the necessary constructs were infiltrated in *N. benthamiana* leaves and harvested at 2 dpi. 1-2 g of tissue was ground in liquid nitrogen, followed by protein extraction, co-immunoprecipitation (co-IP), and Western blotting according to Macho *et al* (2014). The antibodies used are as follows: goat anti-GFP (SICGEN, AB0020-500, 1:5000), mouse anti-RFP (Chromotek, 6G6, 1:5000), goat polyclonal anti-mouse coupled to horseradish peroxidase (Sigma, A2554, 1:15000) and rabbit polyclonal anti-goat coupled to horseradish peroxidase (Sigma, A8919-2ML, 1:20000).

### Yeast two-hybrid (Y2H)

pGBKT7- and pGADT7-derived constructs were co-transformed into the Y2HGold yeast strain (Clontech) using the Frozen-EZ Transformation II kit (Zymo) according to the manufacturer’s instructions. The co-transformants were selected on minimal synthetic defined (SD) media without leucine and tryptophan (double dropout medium, DDO). Interactions were tested on SD media without leucine, tryptophan, histidine, and adenine (quadruple dropout medium, QDO), QDO with X-α-gal (QDO/X) and QDO/X with Aureobasidin A (QDO/X/AbA). Plates were incubated at 30ºC for five to seven days. The pGADT7-T (expressing the SV40 large T-antigen) and pGBKT7-p53 (expressing murine p53) pair were used as a positive control; pGADT7 (AD) and pGBKT7 (BD) empty vectors were used as a negative control.

### Chromatin immunoprecipitation (ChIP)

*A. tumefaciens* clones carrying the binary vector to express RFP-fused proteins were co-infiltrated with those carrying the *pRep::*Rep_WT_ clone in *N. benthamiana* leaves, and tissues were collected at 2 dpi. Chromatin immunoprecipitation (ChIP) assays were performed as described (He *et al*, 2018). In brief, the crosslinking of 2 g of leaf tissue was performed with 1% formaldehyde in 1xPBS buffer and quenched with 0.125 M glycine by vacuum infiltration. Then the tissue was ground to powder and resuspended in Honda buffer (2.5% Ficoll 400, 5% Dextran T40, 0.4 M Sucrose, 25 mM Tris pH 7.4, 10 mM MgCl2, 0.035% b-mercaptoethanol, 1% Protease Inhibitor Cocktail (P9599; Sigma), 1 mM PMSF), homogenized and filtered through Miracloth (Milli-pore). Triton x-100 was added to the supernatant to a final concentration of 0.5%. After spinning at 2000 g for 20 min at 4°C, the pellet was re-suspended in Honda buffer containing 0.1% Triton x-100 and spun at 2000 g for 20 min at 4°C. Isolated nuclei were resuspended in 500 µL of nuclei lysis buffer (0.1% SDS, 50 mm Tris-HCl at pH 8, 10 mm EDTA, 50 µM MG132,1 mM PMSF, 1% Protease Inhibitor Cocktail) and chromatin was sonicated with a Covaris S220 for 3 min (Peak Power: 150, cycles/burst: 200. Duty Factory: 10). Following centrifugation at 16,100 g for 5 min at 4°C, the supernatant was separated and used for input and immunoprecipitation. 300 µl of supernatant was diluted in 900 µl of ChIP dilution buffer, and was pre-cleared with 10 µL of Dynabeads Protein G (Invitrogen) for 1 h at 4°C. After removing the beads, the supernatant was incubated with anti-RFP (Chromotek, 6G6) or anti-IgG antibody (Sigma-Aldrich, NI03) as a negative control overnight at 4°C. The following day, after adding 10 µL of Dynabeads Protein G, the samples were incubated for 2 h at 4°C. Beads were sequentially washed for 5-10 min with 1 mL of the following buffers: two washes with Low-Salt buffer (150 mM NaCl, 0.1% SDS, 1% Triton x-100, 2 mM EDTA, 20 mM Tris pH 8.0), one wash with High-Salt buffer (500 mM NaCl, 0.1% SDS, 1% Triton x-100, 2 mM EDTA, 20 mM Tris pH 8.0), one wash with LiCl buffer (250 mM LiCl, 1% Igepal, 1% sodium deoxycholate, 1 mM EDTA, 10 mM Tris pH 8.0), and two washes with TE buffer (10 mM Tris pH 8.0, 1 mM EDTA). Immunocomplexes were eluted with 150 µL of Elution buffer (1% SDS, 0.1 M NaHCO3) at 65°C for 30 min. After reverse crosslinking overnight, 10 µL of 0.5 M EDTA, 20 µL of 1 M Tris pH 6.5 and 1 µL of proteinase K (Invitrogen) were added to each sample, which was incubated at 45°C for 2 h. DNA was then purified with the QIAquick PCR Purification Kit (Qiagen), and was eluted with 20 µL of ddH2O. IP and input samples were then diluted (1:10 and 1:1000, respectively) and analysed by qPCR. The primers used in this experiment are described in Table S3.

### Statistical analysis

Statistical significance was analyzed with GraphPad Prism 9 or R (v.4.4.1) using ANOVA or Kruskal-Wallis non-parametric test with corresponding post hoc tests (see figure legends for details). Significance levels of *p* < 0.05 (*), *p* < 0.01 (**), *p* < 0.001 (***) and *p* < 0.0001 (****) were chosen. In Figures 1E and S2I different letters represent statistically significant differences (*p* < 0.05). Error bars are shown as mean ± standard deviation or standard error, as indicated in the figure legend.

## ACKNOWLEDGEMENTS

The authors thank Huang Tan, Man Gao, Zhihao Jiang, Clémence Marchal, Shuyi Luo, Nicholay Diaz-Ardila, and Eduardo R. Bejarano for critical reading of the manuscript, and Bettina Stadelhofer and the central facilities at the ZMBP, especially the plant cultivation and the microscopy facilities, for excellent technical support. This work was partially funded by the European Research Council (GemOmics; 101044142) and the DFG (SFB 1101/C08); DMP is the recipient of a Marie Skłodowska-Curie Grant from the European Union’s Horizon Program (VIRALS; 101104619).

