## Supplementary material for "Alternative splicing expands the functional portfolio of a plant virus to control the viral cycle"

**Table S1. Root-mean-square deviation (RMSD) values for splicing-derived Rep isoforms relative to full-length Rep**

**Table S2. Geminiviruses and other CRESS viruses used for splicing predictions**

**Table S3. Primers used in this work**

**Table S4. Plasmids used in this work**

**Figures S1-S6**

**Supplementary references**

**Table S1. Root-mean-square deviation (RMSD) values for splicing-derived Rep isoforms relative to full-length Rep.** RMSD values (Å) were calculated in ChimeraX for each Rep domain. Values in parentheses represent RMSD after structural pruning, with the corresponding number of retained amino acids indicated. RMSD values <1 Å indicate high structural similarity.

| | DNA-binding | Oligomerization | Helicase/ATPase | Total protein |
|---|---|---|---|---|
| Rep289 | 0,614 | - | 9,019 (0,357, 149) | 49,421 (0,172, 89) |
| Rep291 | 0,465 | - | 10,229 (0,326, 150) | 35,272 (0,268, 106) |
| Rep304 | 0,773 | - | 9,547 (0,282, 147) | 34,476 (1,460, 6) |
| Rep306 | 0,488 | - | 12,061 (0,272, 148) | 39,425 (0,298, 154) |
| Rep328 | 3,289 | 2,191 | 10,753 (0,362, 157) | 8,097 (0,646, 238) |

**Table S2. Geminiviruses and other CRESS viruses used for splicing predictions.** AAEO: Africa, Asia, Europe, and Oceania; Sweepoviruses: sweet potato-infecting begomoviruses.

| | Virus name | Family | Genus | Lineage | Reference |
|---|---|---|---|---|---|
| TYLCSV | Tomato yellow leaf curl Sardinia virus | *Geminiviridae* | *Begomovirus* | AAEO region, monopartite | Spain2 L27708 |
| ToLCJV | Tomato leaf curl Joydebpur virus | *Geminiviridae* | *Begomovirus* | AAEO region, monopartite | KM383747 |
| ChiLCV | Chilli leaf curl virus | *Geminiviridae* | *Begomovirus* | AAEO region, monopartite | JN604490.1 |
| ToLCBaV | Tomato leaf curl Bangalore virus | *Geminiviridae* | *Begomovirus* | AAEO region, monopartite | Z48182 |
| TYLCCNV | Tomato yellow leaf curl China virus | *Geminiviridae* | *Begomovirus* | AAEO region, monopartite | OQ753874 |
| AYVV | Ageratum yellow vein virus | *Geminiviridae* | *Begomovirus* | AAEO region, monopartite | KM051527 |
| CLCuGV | Cotton leaf curl Gezira virus | *Geminiviridae* | *Begomovirus* | AAEO region, monopartite | NC_002510.1 |
| ChiLCAV | Chilli leaf curl Ahmedabad virus | *Geminiviridae* | *Begomovirus* | AAEO region, monopartite | KM880103 |
| EACMV | East African cassava mosaic virus | *Geminiviridae* | *Begomovirus* | AAEO region, bipartite | KE2[K201] |
| ACMV | African cassava mosaic virus | *Geminiviridae* | *Begomovirus* | AAEO region, bipartite | MT858793 |
| MYMV | Mungbean yellow mosaic virus | *Geminiviridae* | *Begomovirus* | AAEO region, bipartite | JQ398669.1 |
| ToLCNDV | Tomato leaf curl New Delhi virus | *Geminiviridae* | *Begomovirus* | AAEO region, bipartite | KM977733 |
| ToVCLDeV | Tomato vein clearing leaf deformation virus | *Geminiviridae* | *Begomovirus* | Americas, bipartite | MK423208 |
| ToGLDV | Tomato golden leaf distortion virus | *Geminiviridae* | *Begomovirus* | ? | HM357456 |
| SLCuV | Squash leaf curl virus | *Geminiviridae* | *Begomovirus* | Americas, bipartite | M38183 |
| ToSLCV | Tomato severe leaf curl virus | *Geminiviridae* | *Begomovirus* | Americas, bipartite | AF130415 |
| ToCmMV | Tomato common mosaic virus | *Geminiviridae* | *Begomovirus* | Americas, bipartite | EU710754 |
| CabLCV | Cabbage leaf curl virus | *Geminiviridae* | *Begomovirus* | Americas, bipartite | MH359394 |
| ToGLSV | Tomato golden leaf spot virus | *Geminiviridae* | *Begomovirus* | Americas, bipartite | KC626021 |
| BoGMV | Boerhavia golden mosaic virus | *Geminiviridae* | *Begomovirus* | Americas, bipartite | KY971533 |
| TYMLCV | Tomato yellow margin leaf curl virus | *Geminiviridae* | *Begomovirus* | Americas, bipartite | AY508993 |
| ToLCPVV | Tomato leaf curl purple vein virus | *Geminiviridae* | *Begomovirus* | Americas, bipartite | KY196216 |
| BGMV | Bean golden mosaic virus | *Geminiviridae* | *Begomovirus* | Americas, bipartite | M88686 |
| ToLDV | Tomato leaf distortion virus | *Geminiviridae* | *Begomovirus* | Americas, bipartite | EU710749 |
| SiYNV | Sida common mosaic virus | *Geminiviridae* | *Begomovirus* | Americas, bipartite | EU710751 |
| ToLCV_2 | Tomato interveinal chlorosis virus-2 | *Geminiviridae* | *Begomovirus* | Americas, bipartite | MK087038 |
| ToMoWV | Tomato mottle wrinkle virus | *Geminiviridae* | *Begomovirus* | Americas, bipartite | KM243018 |
| ToLDeV | Tomato leaf deformation virus | *Geminiviridae* | *Begomovirus* | Americas, bipartite | NC014510 |
| AbMV | Abutilon mosaic virus | *Geminiviridae* | *Begomovirus* | Americas, bipartite | X15983 |
| JMV | Jatropha mosaic virus | *Geminiviridae* | *Begomovirus* | Americas, bipartite | KJ174331 |
| CdTAV | Chino del tomate Amazonas virus | *Geminiviridae* | *Begomovirus* | Americas, bipartite | MH243423 |
| ToMoLCV | Tomato mottle leaf curl virus | *Geminiviridae* | *Begomovirus* | Americas, bipartite | BR-BA-11 |

| | | | | | |
|---|---|---|---|---|---|
| PSLDV | Passionfruit severe leaf distortion virus | *Geminiviridae* | *Begomovirus* | Americas, bipartite | FJ972767 |
| ToBYMV | Tomato bright yellow mosaic virus | *Geminiviridae* | *Begomovirus* | Americas, bipartite | KC791690 |
| SPLCLaV | Sweet potato leaf curl Lanzarote virus | *Geminiviridae* | *Begomovirus* | Sweepoviruses | EU839579.2 |
| SPMV | Sweet potato mosaic virus | *Geminiviridae* | *Begomovirus* | Sweepoviruses | FJ969831 |
| SPLCSiV-2 | Sweet potato leaf curl Sichuan virus 2 | *Geminiviridae* | *Begomovirus* | Sweepoviruses | KF156759 |
| SPLCHnV | Sweet potato leaf curl Henan virus | *Geminiviridae* | *Begomovirus* | Sweepoviruses | KC907406 |
| TPCTV | Tomato pseudo-curly top virus | *Geminiviridae* | *Topocuvirus* | - | X84735 |
| OpV1 | Opuntia virus 1 | *Geminiviridae* | *Opunvirus* | - | MN100000 |
| TCTV | Turnip curly top virus | *Geminiviridae* | *Turncurtovirus* | - | GU456685.1 |
| ECSV | Eragrostis curvula streak virus | *Geminiviridae* | *Eragrovirus* | - | FJ665631.1 |
| PNYDV | Pea necrotic yellow dwarf virus | *Nanoviridae* | *Nanovirus* | - | JN133280 |
| PCV2 | Porcine circovirus 2 | *Circoviridae* | *Circovirus* | - | #AJ293868 |

**Table S3. Primers used in this work.**

| Name | Species | Reference | Primer Sequence (5’-3’) |
|---|---|---|---|
| **For cloning:** | | | |
| Rep | Tomato yellow leaf curl virus (TYLCV) | this study | F: ggggacaagtttgtacaaaaaagcaggctccATGCCTCGTTTATTTAAAAT |
| | | | R[1]: ggggaccactttgtacaagaaagctgggtcCGCCTTATTGGTTTCTTCTTG |
| | | | R[2]: ggggaccactttgtacaagaaagctgggtcTTACGCCTTATTGGTTTCTTCTTG |
| | | | F: AGCGTCTCACACCATGCCTCGTTTATTTAAAATATAT |
| | | | R[2]: TACGTCTCTCCTTTTACGCCTTATTGGTTTCTTCTTGGCTATC |
| Rep$_{289}$ | TYLCV | this study | F: ACTCTTCAAGTTCATCTCCTCTAGCTGATC |
| | | | R:GATCAGCTAGAGGAGATGAACTTGAAGAGT |
| Rep$_{291}$ | TYLCV | this study | F: CAAGTTCATCTGGAACTCCTCTAGCTGATC |
| | | | R: GATCAGCTAGAGGAGTTCCAGATGAACTTG |
| Rep$_{304}$ | TYLCV | this study | F: GCACTCAATTCAGATGAACTTGAAGA |
| | | | R: TCTTCAAGTTCATCTGAATTGAGTGC |
| Rep$_{306}$ | TYLCV | this study | F: GCACTCAATTCAGTTCCAGATGAACT |
| | | | R: AGTTCATCTGGAACTGAATTGAGTGC |
| Rep$_{328}$ | TYLCV | this study | F: AATGTTCGGATGGAAATGTGGTTCCCCATT |
| | | | R: AATGGGGAACCACATTTCCATCCGAACATT |
| Rep promoter / TYLCV intergenic region | TYLCV | this study | F: ggggacaagtttgtacaaaaaagcaggctccATTGCAAGACAAAATACTTGGGGAC |
| | | | R: ggggaccactttgtacaagaaagctgggtcGTTGAAATGAATCGGTGTCCCTC |
| Rep promoter-Rep-C2-C3 | TYLCV | this study | R: ggggaccactttgtacaagaaagctgggtcTTAATAAAATTTATATTTTATATCA |
| **For site directed mutagenesis:** | | | |
| Rep$_{\#5\text{-}8\Delta d,as}$, donor sites #5-6 | TYLCV | this study | F: ATGCCGAAGCACTCAATTCCGGAAATAAATCCGAGGCCCT |
| | | | R: AGGGCCTCGGATTTATTTCCGGAATTGAGTGCTTCGGCAT |
| Rep$_{\#5\text{-}8\Delta d,as}$, donor sites #7-8 | TYLCV | this study | F: GCAGATCAGCTAGAGGAGGCCAGCAATCTGCCAACGACGC |
| | | | R: GCGTCGTTGGCAGATTGCTGGCCTCCTCTAGCTGATCTGC |

| | | | |
|---|---|---|---|
| **For qPCR:** | | | |
| Rep | TYLCV | Wang *et al*, (2017) | F: TGAGAACGTCGTGTCTTCCG |
| | | | R: TGACGTTGTACCACGCATCA |
| C2 | TYLCV | Wang *et al*, (2017) | F: ACCTTCGTCACCCTCTACGA |
| | | | R: TGACGTTGTACCACGCATCA |
| C3 | TYLCV | Wang *et al*, (2017) | F: TGAGAACGTCGTGTCTTCCG |
| | | | R: ACAATACATGATCAACTGCTCTGA |
| CP | TYLCV | Wang *et al* (2022) | F: TGGAAGCAGCCCAATGGATT |
| | | | R: GTTCTCGTACTTGGCTGCCT |
| *Internal transcript spacer 25SrDNA (ITS)* | *Nicotiana benthamiana* | Mason *et al* (2008) | F: ATAACCGCATCAGGTCTCCA |
| | | | R: CCGAAGTTACGGATCCATTT |
| *NbEF1α (Elongation factor 1-alpha)* | *Nicotiana benthamiana* | Segonzac *et al* (2011) | F: AAGGTCCAGTATGCCTGGGTGCTTGAC |
| | | | R: AAGAATTCACAGGGACAGTTCCAATACCAC |
| q2IR-GFP | TYLCV | this study | F: TGTTCCATGGCCAACACTTG |
| | | | R: ACGTGTCTTGTAGTTCCCGT |
| qIR-ChIP | TYLCV | this study | F: CCAAATAGCCATTAGGTGTCCAG |
| | | | R: GAATCGGTGTCCCTCAAAGC |
| *GFP* | N/A | | F: TGCTGCTGCCCGACAACCACTAC |
| | | | R: CTTGTACAGCTCGTCCATGCC |
| For RT-PCR: | | | |
| Rep | TYLCV | (Wang *et al*, 2017, 2022) | F: ATCCGAACATTCAGGCAGCT |
| | | | R:TGACGTTGTACCACGCATCA |
| Rep splicing event #5 | TYLCV | Pott *et al* (2025) | F: GACCCACTCTTCAAGTTCATCTGAAT |
| | | | R: CAACGGTTCTTCGACCTGGT |
| Rep | EACMV | this study | F: CATGGTCTTCCCTGTACGACT |
| | | | R: TCGTCCGATGTCAAGGCTTA |
| Rep | PNYDV | this study | F: TGCTGGTATGAAGAGGCTCA |
| | | | R: ACACTTTGGTATTTGCCTGATTG |
| *NbActin* | *Nicotiana benthamiana* | Pott *et al* (2025) | F: ATGTTCACCACCTCAGCTGA |
| | | | R: GGGAAGCCAAGATAGAGC |

F: forward; R: reverse. In lowercase, attB sites for Gateway compatible cloning. F primer includes attB1 sequence; R primer includes attB2 sequence. Superscripts indicate without stop codon (1) or with stop codon (2). Underlined sequences correspond to *Esp31* sites and corresponding overhangs for Golden Gate compatible cloning.

**Table S4. Plasmids used in this work**

| Construct name | Source | Resistance | Destination vector name (source) |
|---|---|---|---|
| TYLCV | Rosas-Diaz *et al* (2018) | Spectinomycin | pGWB501; RRID:Addgene_74843 |
| TYLCV_C4$_{1\text{-}8}$ | Rosas-Diaz *et al* (2018) | Spectinomycin | pGWB501; RRID:Addgene_74843 |
| TYLCV$_{Rep\#5\text{-}8\Delta as}$ | Pott *et al* (2025) | Spectinomycin | pGWB501; RRID:Addgene_74843 |
| TYLCV_C4 $_{1\text{-}8\ Rep\#5\text{-}8\Delta d,as}$ | this study | Spectinomycin | pGWB501; RRID:Addgene_74843 |
| 35S:GFP | Medina-Puche *et al* (2020) | Kanamycin | pGWB2; NCBI:txid419546; AB289765 |
| 35S:Rep | Pott *et al* (2025) | Spectinomycin | pGWB502; RRID:Addgene_74844 |
| 35S:Rep-GFP | this study | Spectinomycin | pGWB505; RRID:Addgene_74847 |
| 35S:Rep-RFP | this study | Spectinomycin | pGWB554; RRID:Addgene_74884 |
| 35S:Rep$_{289}$ | this study | Spectinomycin | pGWB502; RRID:Addgene_74844 |
| 35S:Rep$_{289}$-GFP | this study | Spectinomycin | pGWB505; RRID:Addgene_74847 |
| 35S:Rep$_{289}$-RFP | this study | Spectinomycin | pGWB554; RRID:Addgene_74884 |
| 35S:Rep$_{291}$ | this study | Spectinomycin | pGWB502; RRID:Addgene_74844 |
| 35S:Rep$_{291}$-GFP | this study | Spectinomycin | pGWB505; RRID:Addgene_74847 |
| 35S:Rep$_{291}$-RFP | this study | Spectinomycin | pGWB554; RRID:Addgene_74884 |
| 35S:Rep$_{304}$ | this study | Spectinomycin | pGWB502; RRID:Addgene_74844 |
| 35S:Rep$_{304}$-GFP | this study | Spectinomycin | pGWB505; RRID:Addgene_74847 |
| 35S:Rep$_{304}$-RFP | this study | Spectinomycin | pGWB554; RRID:Addgene_74884 |
| 35S:Rep$_{306}$ | this study | Spectinomycin | pGWB502; RRID:Addgene_74844 |
| 35S:Rep$_{306}$-GFP | this study | Spectinomycin | pGWB505; RRID:Addgene_74847 |
| 35S:Rep$_{306}$-RFP | this study | Spectinomycin | pGWB554; RRID:Addgene_74884 |
| 35S:Rep$_{328}$ | this study | Spectinomycin | pGWB502; RRID:Addgene_74844 |
| 35S:Rep$_{328}$-GFP | this study | Spectinomycin | pGWB505; RRID:Addgene_74847 |
| 35S:Rep$_{328}$-RFP | this study | Spectinomycin | pGWB554; RRID:Addgene_74884 |
| 35S:Rep$_{\#5\text{-}8\Delta as}$ | Pott *et al* (2025) | Spectinomycin | pGWB554; RRID:Addgene_74884 |
| 35S:Rep$_{\#5\text{-}8\Delta d,as}$ | this study | Spectinomycin | pGWB554; RRID:Addgene_74884 |
| pRep:GFP | this study | Spectinomycin | pGB504; RRID:Addgene_74846 |

| | | | |
|---|---|---|---|
| pRep-Rep | this study | Spectinomycin | pGWB501; RRID:Addgene_74843 |
| pRep:Rep$_{null}$-C2-C3 | this study | Spectinomycin | pGWB501; RRID:Addgene_74843 |
| AD-Rep | this study | Ampicillin | pGTT2t7AD |
| AD-Rep$_{291}$ | this study | Ampicillin | pGADT7-GW; RRID:Addgene_61702 |
| AD-Rep$_{306}$ | this study | Ampicillin | pGADT7-GW; RRID:Addgene_61702 |
| BD-Rep | this study | Kanamycin | pGTT2t7BK |
| BD-Rep$_{291}$ | this study | Kanamycin | pGBKT7-GW; RRID:Addgene_61702 |
| BD-Rep$_{306}$ | this study | Kanamycin | pGBKT7-GW; RRID:Addgene_61702 |
| PVX$_{C4}$ | this study | Kanamycin | pICH31160 (Icon Genetics) |

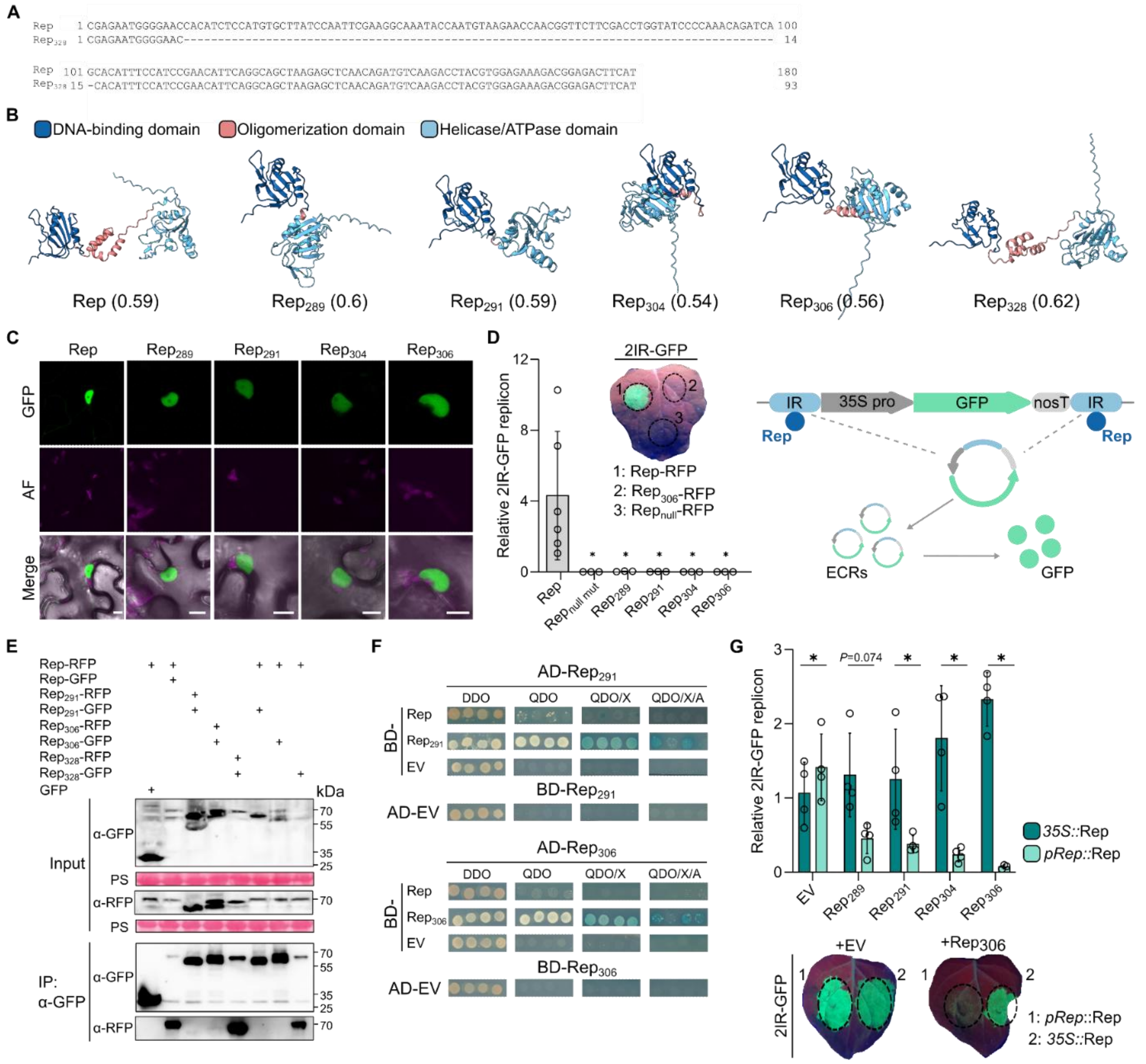


**Figure S1. Characterization of Rep spliced isoforms.** (A) Confirmation by Sanger sequencing of an additional splicing event in Rep with an intron of 87 bp, and alignment with Rep full length. This event leads to $Rep_{328}$, lacking 29 amino acids (position 56 to 84) in the N-terminal DNA binding domain.

(B) Alphafold3 prediction (https://alphafoldserver.com/, Abramson *et al* (2024)) of TYLCV Rep and Rep spliced isoforms. Spliced isoform conformation is shown after alignment with Rep full length in ChimeraX. Predicted template modelling score (pTM) are indicated in parentheses.

(C) Subcellular localization of Rep and Rep splicing-derived variants fused to C-ter GFP and expressed in *Nicotiana benthamiana*. Scale bar = 10 µm.

(D) Rep spliced variants do not support GFP replicon accumulation in the virus-free reporter system 2IR-GFP. A representative leaf exposed to UV light is shown, in which Rep, Rep null mutant and $Rep_{306}$ have been infiltrated. GFP replicons were measured by qPCR at 2 dpi.

Error bars represent SD. The experiment was repeated three times with similar results (n=3), and leaf disks expressing the same construct were then pooled together for qPCR. A schematic representation of the 2IR-GFP cassette is shown. IR: TYLCV intergenic region; 35S pro: 35S cauliflower mosaic virus (CaMV) promoter; NosT: NOS terminator. Upon Rep expression in *N. benthamiana* 2IR-GFP plants, the viral Rep specifically binds the IR, triggering the generation of GFP extra-chromosomal replicons (ECRs), which results in an overexpression of the *GFP* transgene and accumulation of the fluorescent protein.

(E) Rep spliced variants ($Rep_{306}$ and $Rep_{291}$) weakly interact with themselves (lanes 3 and 4, bottom blot showing α-RFP signal in the immunoprecipitated samples), but not with unspliced Rep by co-immunoprecipitation (co-IP). Rep and $Rep_{328}$ homo- and heterodimerization are used as positive controls. $Rep_{306}$ and $Rep_{291}$ self-interaction was tested three times with similar results; $Rep_{306}$ and $Rep_{291}$ interaction with full-length Rep was tested twice.

(F) Rep spliced variants ($Rep_{306}$ and $Rep_{291}$) can interact with themselves, but not with unspliced Rep by yeast two hybrid (Y2H). The interactions can be observed on QDO, QDO/X and QDO/X/A plates, the latter showing high-stringency interactions. Empty AD/BD vectors (AD-EV and BD-EV, respectively) were used as negative controls. AD: GAL4 activation domain; BD: DNA-binding domain. $Rep_{306}$ and $Rep_{291}$ self-interaction were tested three times with similar results; $Rep_{306}$ and $Rep_{291}$ interaction with full-length Rep were tested twice.

(G) Effect of Rep spliced variants on GFP replicon accumulation when *35S::*Rep or *pRep::*Rep is expressed in 2IR-GFP *N. benthamiana* plants. The splicing-derived isoforms and either *35S::*Rep or *pRep::*Rep were co-inoculated at the same time. GFP replicons were measured by qPCR at 2 dpi. Representative leaves exposed to UV light are shown, in which *35S::*Rep (right half of the leaf) and *pRep::*Rep (left half) are expressed, either with an empty vector as a control or $Rep_{306}$-RFP. Error bars represent SD. This experiment was repeated three times with similar results (n=3-4). Results from one experiment are shown. A nonparametric two-way ANOVA test using the aligned rank transform was performed, followed by Tukey post hoc tests to show statistically significant differences.

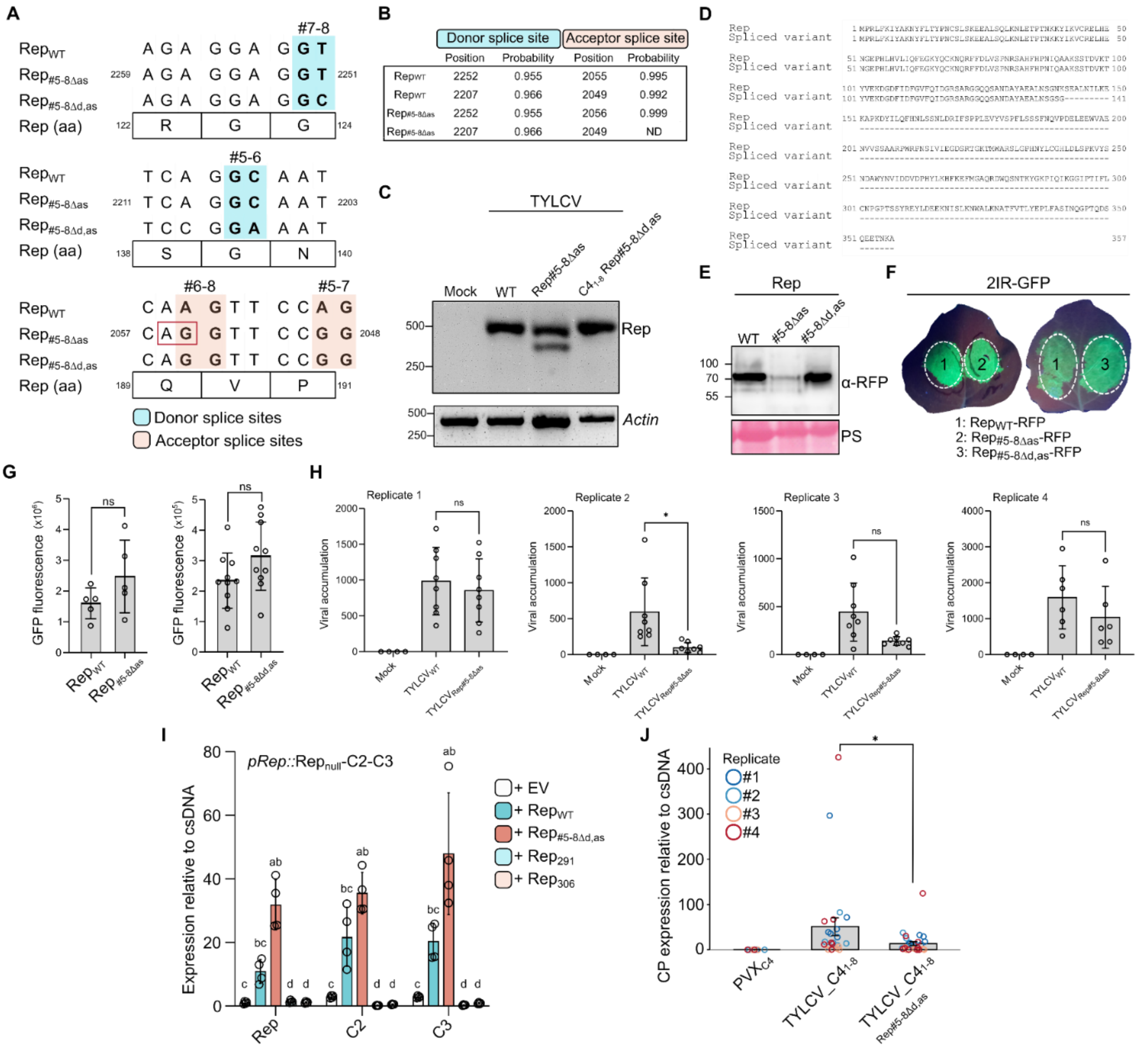

**Figure S2. Characterization of Rep splicing mutants Rep#5-8Δas and Rep#5-8Δd,as.** (A) To generate Rep#5-8Δd,as, point mutations were introduced in the 5' (donor) splice sites for splicing events #5-8 (highlighted in blue), without affecting the Rep amino acid sequence. Positions of the mutations (in nt) in the TYLCV infectious clone (AJ489258) are shown. Mutations in the 3' (acceptor) splice sites for Rep#5-8Δas are also present in the Rep#5-8Δd,as mutant and are shown in orange in the bottom alignment.

(B) A novel acceptor splice site (shown in the red square in A, alignment at the bottom) generated by mutating acceptor splice sites #6 and #8 is located 1 bp upstream of the canonical acceptor splice site and is predicted with very high probability by the predictor tool Spliceator (Scalzitti *et al*, 2021). ND: not detected.

(C) This novel acceptor splice site leads to the generation of a spliced isoform in Rep#5-8Δas, similar to splicing events #6 and #8 (with a 1-bp shift) and can be observed on an agarose gel showing cDNA from TYLCVRep#5-8Δas-expressing samples. The primers amplify a 516-bp fragment of Rep transcript, flanking splicing events #5-8. A shorter transcript corresponding to the novel spliced isoform is present in $TYLCV_{Rep\#5\text{-}8\Delta as}$ but not in $TYCLV_{WT}$ or $TYLCV\_C4_{1\text{-}8\ Rep\#5\text{-}8\Delta d,as}$

(D) Amino acid sequence of the spliced isoform generated in $Rep_{\#5\text{-}8\Delta as}$. The sequence is aligned with $Rep_{WT}$ amino acid sequence.

(E) Western blot showing the accumulation of Rep-, $Rep_{\#5\text{-}8\Delta as\text{-}}$ and $Rep_{\#5\text{-}8\Delta d,as}$-RFP when transiently expressed in *N. benthamiana* leaves. Samples were harvested at 2 dpi.

(F) Rep splicing mutants are not impaired in their ability to initiate GFP replicon accumulation in 2IR-GFP *N. benthamiana* leaves. Representative leaves under UV light are shown, in which Rep (left half of the leaf) and Rep splicing mutants (right half) were infiltrated.

(G) GFP fluorescence measured in a plate reader in disc leaves from (F). Error bars represent SD. This experiment was repeated three times with similar results (n=5-10). Results from one experiment are shown. Statistical significance was assessed using two-tailed Student's t-tests.

(H) Accumulation of TYLCV and $TYLCV_{Rep\#5\text{-}8\Delta as}$ in local infection assays in *N. benthamiana* (four independent replicates, n=6-8). Viral accumulation was measured by qPCR at 3 dpi. Replicate 1 is shown in Figure 2C. Error bars represent SD. Statistical analyses were performed separately for each replicate: replicates 2 and 3 were analyzed using a Kruskal–Wallis test followed by Dunn's post hoc test, replicate 1 using Dunnett T3 post hoc test, and replicate 4 using Tukey's post hoc test, to evaluate significant differences between TYLCV and TYLCV.

(I) Rep, C2 and C3 expression from a partial genome comprising a fragment from the intergenic region (containing the Rep promoter) to the end of the C3, in which Rep has been mutated to render it unable to produce the corresponding protein (*pRep::*$Rep_{null}$-C2-C3). Expression in *trans* of Rep splicing-derived variants (represented by $Rep_{291}$ and $Rep_{306}$) repress the expression of Rep, C2, and C3, while $Rep_{\#5\text{-}8\Delta d,as}$ increases their expression compared to $Rep_{WT}$. Error bars represent SD. This experiment was repeated three times with similar results (n=4). Results from one experiment are shown. Different letters above the bars indicate statistically significant differences at $p < 0.05$, after a nonparametric two-way ANOVA test using the aligned rank transform followed by Tukey-adjusted post hoc tests.

(J) CP expression in samples systemically infected with $TYLCV\_C4_{1\text{-}8}$ or $TYLCV\_C4_{1\text{-}8\ Rep\#5\text{-}8\Delta d,as}$, co-inoculated with $PVX_{C4}$ (14 dpi). Expression is normalized to TYLCV csDNA. Error bars represent SE. This experiment was repeated four times with similar results (n=6-8). A linear mixed-effects model with the genotypes as a fixed effect and the replicates as a random effect was used. Asterisks indicate statistically significant differences after pairwise comparisons among groups using estimated marginal means with Tukey-adjusted p-values.

**A**

```
RepEACMV (326-687pb)   1 AATTTCTTGACGATGGAATTTTCCAAGTCGATGCCCGAAGTGCCAGGGGGGAGGGCCAGCATTTAGCTCAGGTATATGCAGACGCGTTGAATGCTTCTTC 100
RepEACMV lower_band    1 AATTTCTTGACGATGGAATTTTCCAAGTCGATGCCCGAAGTGCCAGGGGGGAGGGCCAGCATTTAGCTCAG--------------------------------  68

RepEACMV (326-687pb) 101 TAAATCAGAAGCTCTTCAAATTATCAAAGAAAAAGATCCCAAGTCTTTTTTTTTACAGTTCCATAACATATCTGCTAACGCAGATCGAATCTTCCAGGCT 200
RepEACMV lower_band   69 -----------------------------------------------------------------------------------ATCGAATCTTCCAGGCT  88

RepEACMV (326-687pb) 201 CCGCCCCAAACTTACGTTAGTCCGTTCTTATCATCCTCTTTTACTAACGTTCCTGAGGAACTTGAAGTCTGGGTTTCCGAGAACGTGATGGGTTCCGCTG 300
RepEACMV lower_band   89 CCGCCCCAAACTTACGTTAGTCCGTTCTTATCATCCTCTTTTACTAACGTTCCTGAGGAACTTGAAGTCTGGGTTTCCGAGAACGTGATGGGTTCCGCTG 188

RepEACMV (326-687pb) 301 CGCGGCCTTGGAGACCCAACAGTATTGTTCTAGAAGGTGATAGTCGTACAGGGAAGACCATG                                      362
RepEACMV lower_band  189 CGCGGCCTTGGAGACCCAACAGTATTGTTCTAGAAGGTGATAGTCGTACAGGGAAGACCATG                                      250
```

**B**

```
RepEACMV                    1 MPRAGRFQINAKNYFITYPRCSLTKEEALSQLKAFSYPTNIKFIRVCRELHQDGVPHLHVLIQFEGKFQCTNPRFFDLISPSRSTHFHPNIQGAKSSSDV 100
RepEACMV spliced isoform    1 MPRAGRFQINAKNYFITYPRCSLTKEEALSQLKAFSYPTNIKFIRVCRELHQDGVPHLHVLIQFEGKFQCTNPRFFDLISPSRSTHFHPNIQGAKSSSDV 100

RepEACMV                  101 KAYIEKGGEFLDDGIFQVDARSARGEGQHLAQVYADALNASSKSEALQIIKEKDPKSFFLQFHNISANADRIFQAPPQTYVSPFLSSSFTNVPEELEVWV 200
RepEACMV spliced isoform  101 KAYIEKGGEFLDDGIFQVDARSARGEGQHLAQIESSRLR-----------------------------------PKLTLVRSY----------------- 148

RepEACMV                  201 SENVMGSAARPWRPNSIVLEGDSRTGKTMWARSLGPHNYLCGHLDLSPKVYSNDAWYNVIDDVDPHYLKHFKEFMGAQRDWQSNTKYGKPIQIKGGIPTI 300
RepEACMV spliced isoform  149 --------------------------------------------------------------HPLLLTFLRN--------------------LKSGFP-- 164

RepEACMV                  301 FLCNPGPTSSYKEFLDEEKNQSLKAWALKNATFVTLHEPLFSSAHQSPTPHSEDQGHQT                                         359
RepEACMV spliced isoform  165 ---------------------------------------------------------RT                                         166
```

**Figure S3**. **Confirmation of a splicing event in the Rep transcript from EACMV by Sanger sequencing.** (A) Nucleotide alignment between the Rep sequence from EACMV (full length) and the sequenced spliced isoform detected in cDNA from *N. benthamiana* leaves expressing *35S::*Rep$_{EACMV.}$

(B) Alignment between Rep from EACMV amino acid sequence and the generated Rep spliced isoform.

**A**

```
RepPNYDV (205-761pb)      1 AGGAGAGGTACACAAGGAGAAGCTCGTGCTTATGCTATGAAGGAAGACTCCCGTCTTGAAGGTCCATGGGAAGAAGGTGAGTTTCACCTCACAGTGGAAG 100
RepPNYDV lower band       1 AGGAGAGGTACACAAGGAGAAGCTCGTGCTTATGCTATGAAGGAAGACTCCCGTCTTGAAGGTCCATGGGAAGAAGGTGAGTTTCACCTCACAGTGGAAG 100

RepPNYDV_(205-761pb)    101 ACAAGCTTCGTGAAGTTATGGAAGATATGAAGAACACAGGTAAACGTCCTATTGAATACATAGAAGAGTGTTGTAACACCTACGACAAGTCGTCCGGGAC 200
RepPNYDV lower band     101 ACAAGCTTCGTGAA-------------------------------------------------------------------------------------- 114

RepPNYDV_(205-761pb)    201 TCTCAGGGAATACCAAGGTGAGTTACGGAAGAAACGTGCCATTAATGGATGGCAGTTGCAGAGGAAGCCATGGATGGACGAGGTCGAAAGCTTGCTTGAG 300
RepPNYDV lower band         ----------------------------------------------------------------------------------------------------

RepPNYDV_(205-761pb)    301 ACCAGAGATGGAAGAAGAATCATATGGGTGTATGGACCACAAGGTGGGGAAGGGAAAACATCTTTCGCAAAGCATCTTGTAAAGACGCGTGATGCTTTCT 400
RepPNYDV lower band         ----------------------------------------------------------------------------------------------------

RepPNYDV_(205-761pb)    401 ATTCCACTGGCGGAAAGACAGCCGACATTGCTTTCGCTTGGGACCACCAACCAATAGTGCTTTTCGACTTTCCTCGAAGCTTCGAGGAATATGTTAACTA 500
RepPNYDV lower band         ----------------------------------------------------------------------------------------------------

RepPNYDV_(205-761pb)    501 TGGTGTTATGGAACAATTAAAGAATGGGATTATTCAATCAGGCAAATACCAAAGTGT                                            557
RepPNYDV lower band     115 ----GTTATGGAACAATTAAAGAATGGGATTATTCAATCAGGCAAATACCAAAGTGT                                            167
```

**B**

```
RepPNYDV                    1 MSRQVICWCFTLNNPLSLLSLHESMKYLVYQREQGESGNIHFQGYVEMKKRTSLAGMKRLIPGAHFEKRRGTQGEARAYAMKEDSRLEGPWEEGEFHLTV 100
RepPNYDV spliced isoform    1 MSRQVICWCFTLNNPLSLLSLHESMKYLVYQREQGESGNIHFQGYVEMKKRTSLAGMKRLIPGAHFEKRRGTQGEARAYAMKEDSRLEGPWEEGEFHLTV 100

RepPNYDV                  101 EDKLREVMEDMKNTGKRPIEYIEECCNTYDKSSGTLREYQGELRKKRAINGWQLQRKPWMDEVESLLETRDGRRIIWVYGPQGGEGKTSFAKHLVKTRDA 200
RepPNYDV spliced isoform  101 EDKLRE---------------------------------------------------------------------------------------------- 106

RepPNYDV                  201 FYSTGGKTADIAFAWDHQPIVLFDFPRSFEEYVNYGVMEQLKNGIIQSGKYQSVIKYAEYVEVIAFANFTPRSGMFSEDRIIIVNV              286
RepPNYDV spliced isoform  107 ------------------------------------VMEQLKNGIIQSGKYQSVIKYAEYVEVIAFANFTPRSGMFSEDRIIIVNV              156
```

**Figure S4**. **Confirmation of a splicing event in the Rep transcript from PNYDV by Sanger sequencing.** (A) Nucleotide alignment between the Rep sequence from PNYDV (full length) and the sequenced spliced isoform detected in cDNA from *N. benthamiana* leaves expressing *35S::*$Rep_{PNYDV.}$

(B) Alignment between Rep from PNYDV amino acid sequence and the generated Rep spliced isoform.

DNA-binding domain
Oligomerization domain
Helicase/ATPase domain

```
              1          10        20        30          40             50
 1 TYLCV     ...M.PRLFKIYAKNYFLTYPNCSLSKEEALSQLKNLE.TPTNKKYIKV.....CREL E
 2 ToLDeV    .MPP.PKAFKINAKNYFLTYPHCNLTKEEALSQLQNLE.TPTNKKYIKI.....AREL E
 3 ToLCV-2   .MPP.PKRFQIKAKNYFLTYPQCSLSKEEALSQLQALS.TPTNKKFIKI.....CREL E
 4 ToMoWV    .MPL.PKRFKISSKNYFITYPKCSLTKEDALSQLQALQ.TPTNKKFIKI.....CREL E
 5 ToLDV     .MPP.PKRFKISSKNYFVTYPHCSLTKEEALAQLKLLN.TPTNKKYIKI.....CREL E
 6 BGMV      .MPP.PKRFKINAKNYFLTYPQCSITKESAIEQLQNLQ.TPVNKKYIRI.....CREI E
 7 SiYNV     .MPR.KGSFSIKAKNYFLTYPQCSLPKEEALEQIRVIQ.TPVNKKFIKV.....AREL E
 8 ToLCPVV   .MPP.PKRFRISSKNYFLTYPKCSISKEEALSQLQALE.TPVNKKYIRV.....CREL D
 9 ToGLSV    .MGR..............SSKPDVHQESA.......................RRGT W
10 SLCuV     NMPRNPNSFRLTARNIFLTYPRCDVPKEEVLEMLLHLSWSVVKPNYVRV.....AREE S
11 CabLCV    .MPRNPKSFRLAARNIFLTYPQCDIPKDEALQMLQTLSWSVVKPTYIRV.....AREE S
12 ToCmMV    .MPRNPNIFRLTARNIFLTYPQCDIPKDEALQMLQSLPWSVVKPTYIRV.....AREE S
13 ToSLCV    .MPRNPNLFRLAAKNIFLTYPQCDIPKDEAIEMLQNLPWSVVKPTYIRV.....AREE A
14 TYMLCV    .MPT.ARAFKINAKNYFLTYPKCSLSKEEALSQLQNLI.TPTNKKFIKV.....CKEL E
15 BoGMV     .MPR.KGAFSINARNYFLTYPKCSLTKEEALSQIQKLN.TPVNKKFIKI.....CREL E
16 CdTAV     .MPP.PKRFKISSKNYFLTYPHCSLTKDDALSQLINLQ.TPTNKLFIRV.....AREF E
17 JMV       .MPS.VRRFKVSAKNYFLTYPQCSLSKEEALSQLQNLE.TPVNKKFIKI.....CREL E
18 AbMV      .MPP.PKKFRVQAKNYFLTYPQCSLTKDEALSQLQNLN.TPVNKKFIKI.....CREL E
19 ToVCLDeV  .MPSAPKRFRVQSKNFFLTYPHCSLSKEEALSQLQALG.TPVNKKFIRV.....CREL E
20 PSLDV     .MPSEPRRFRVNCRNFFLTYPKCSLSKEEALSQLLALE.TPTNKKFIRV.....SREL E
21 ToMoLCV   .MPL.PRRFRIHSKNYFLTYPKCSLTKEEALSQLLGLQ.TPVNKLFVRV.....AREL E
22 ToGLDV    .MPP.PKRFKINSKNYFLTYPQCSLTKEEALYQLQNLN.TPTNKLFIRV.....AREL D
23 SPLCSiV-2 .MAA.PKKFQINSKNYFLTYPRCSLSKEEALQQLLNIQ.TPTNKKYIHI.....AREL E
24 SPLCHnV   .MVR.APGFRLQAKNIFLTYPKCPLSKDTVLELLKGIQ.CPSDKLFIRV.....AQEK Q
25 SPMV      .MAP.PKRFKIQAKNYFLTYPHCSLSKEEVLDQLKKIQ.TPVNKKYIHI.....AREL E
26 SPLCLaV   .MPR.AGRFNIKAKNYFLTYPQCSLTKEEALDQLLHLN.TPTNKKFIKI.....CREL E
27 MYMV      .MPR.LGRFAINAKNYFLTYPRCPLTKEDALQQLLALS.TPVNKKFIRV.....CREL E
28 EACMV     .MPR.AGRFQINAKNYFITYPRCSLTKEEALSQLKAFS.YPTNIKFIRV.....CREL Q
29 CLCuGV    .MAP.PHRFQIYAKNYFLTFPKCSLTKEEALEQIQKIS.TASNKKYIKI.....CREL E
30 ACMV      ..MR.TPRFRVQAKNVFLTYPKCSIPKEHLLSFIQTLS.LPSNPKFIKI.....CREL Q
31 ToLCNDV   .MAP.PSRFRINAKNYFLTYPKCSLTKEEALSQLQTLE.TPTSKKFIKI.....CREL E
32 AYVV      .MAP.PKRFLINCKNYFLTYPQCSLTKEEALSQLQNLN.TPTNKKYIKICSPGDCREL E
33 TYLCSV    .MAQ.PKRFQINAKHYFLTFPKCSLSKEEALEQLLQLQ.TPTNKKYIKI.....CREL E
34 TYLCCNV   .MAP.PNRFRINAKNYFLTYPHCSLSKEEALSQIQTLE.TPVNKLFIRI.....CREL E
35 ToLCBaV   .MPR.PGRFNINAKNYFLTYPKCSLSKERHLSQLQTVK.TPTSKLFIRV.....CREL V
36 ToLCJV    .MAP.PNKFRINAKNYFLTYPKCSLTKEEALSQLQNLE.TPVNKLFIRV.....CREL E
37 ChiLCAV   .MPR.TGRFNINAKNYFLTYPNCSLTKEEALSQLQNLQ.TPVNKLFIRV.....CREL E
38 ChiLCV    .MPR.AGRFNINAKNYFLTYPNCSLTKEEALSQLQNLE.TPVNKLFIRV.....CREL E

                    60         70           80         90         100
 1 TYLCV     NGEPHLHV I FEGKYQCKNQRF .DLVSPNRSAHFHPNIQAAKSSTDV.KTYVEKDGDF
 2 ToLDeV    DGQHHLHV L FQGKFQCKNNRF .DLVSPTRSTHFHPNVQGAKSSTDV.NAYVDKDGDT
 3 ToLCV-2   DGQPHLHV L FEGNFCCQNQRF .DLVSPTRSAHFHPNIQRAKSSSDV.KSYVDKDGDT
 4 ToMoWV    NGEPHLHV I FEGKYQCTNNRF .DLVSPTRSTHFHPNIQGAKSSSDV.KSYIDKDGDT
 5 ToLDV     NGEPHLHV I FEGKYQCTNNRF .DLVSPSRSAHFHPNIQGAKSSSDV.KSYIDKDGDT
 6 BGMV      NGEPHLHA I FEGKFQCTNCRV .DLKHPTTSSVSHANIQSAKSSSDV.KSYIEKDGDY
 7 SiYNV     DGQPHLHV V FEGKFNCTNSRL .DLVSPSRSTHFHPNIQGAKSSSDV.KSYVEKDGDT
 8 ToLCPVV   NGEPHLHA L FEAKFVCTNCRF .DLRHPSNSNVCHGKYESCKSSSDA.KSYIEKDGDY
 9 ToGLSV    TANLIFTA I LIRGVEHQGCSI RPLLTPRRSGKFSPKRPGSQIRQRVHESDSPRDSLY
10 SLCuV     DGSPHLHC I LSGKSNIKDAGF .DLTHPRRSARFHPNIQAAKDTNAV.KNYITKEGDY
11 CabLCV    DGFPHLHC I LSGKSNIKDARF .DITHPRRSANFHPNIQAAKDTNAV.KNYITKDGDY
12 ToCmMV    DGFPHLHC V LSGKSNIKDVRF .DLTHPRRSTGFHPNVQAAKDTNAV.KNYITKEGDY
13 ToSLCV    DGFPHLHC I LSGKSNIKDARF .DLTHPRRSANFHPNAQAAKDTNAV.QNYITKEGDY
14 TYMLCV    DGEPHLHV I FEGKYQCKNNRF .DLVSPTRSTHFHPNIQGAKSSSDV.KSYIDKDGDT
15 BoGMV     DGQPHLHV I FEGKYNCSNNRF .DLVSPTRSAHFHPNIQGAKSSSDV.KSYIAKDGDT
16 CdTAV     NGEPHLHV I FEGKYQCTNNRF .DLVSPTRSAHFHPNIQGAKSSSDV.KAYMEKDGDF
17 JMV       NGEPHLHV I FEGKYQCTNHRF .DLVSPSRSAHFHPNIQGAKSSSDV.KSYIDKDGDT
18 AbMV      NGEPHLHV I FEGKYQCTNNRF .DLVSPTRSAHFHPNIQGAKSSSDV.KSYIDKDGDT
19 ToVCLDeV  NGEPHLHV I FEGKYVCTNQRF .DLVSPTRSAHFHPNIQGAKSSSDV.KTYVEKHGDF
20 PSLDV     DGQPHLHV I FEGKYNCQNNRF .DLVSTTRSTHFHPNIQGAKSSSDV.KTYVEKDGDF
21 ToMoLCV   DGEPHLHV I FEGKLQCTNERL .DLISPTRSAHFHPNVQGAKSSSDV.KSYVEKDGDF
22 ToGLDV    DGEPHLHV I FEGKYQCTNHRF .DLVSPTRSAHFHPNIQGAKEKLRCYGIRGERRRLH
23 SPLCSiV-2 NGEPHLHM V FEGKFRCTNQRF .DLVSPTRSAHFHPNIQGAKSSSDV.KSYVEKDGDT
24 SPLCHnV   DGSLHLHA I FKGKCQFTNPRH .DLTHPNSSTQYHPNFQSAKSSSDV.KAYIEKDQDF
25 SPMV      DGQPHIHV L FEGKFVCTNQRL .DLVSPTRSAHFHPNIQGAKSSSDV.KSYVDKDGDT
26 SPLCLaV   NGEPHLHV L FEGNYQCTNQRF .DLVSPSRSSHFHPNIQRAKSSSDV.KSYVDKDGDT
27 MYMV      DGEPHLHV L FEGKLQTKNERF .DLVSTTRSAHYHPNVQAAKSASDV.KSYMDKDGDV
28 EACMV     DGVPHLHV I FEGKFQCTNPRF .DLISPSRSTHFHPNIQGAKSSSDV.KAYIEKGGEF
29 CLCuGV    DGQPHLHV L FEGKFKCQNQRL .DLVSPNRSAHFHPNIQGAKSSSDV.KSYIDKDGDT
30 ACMV      NGEPHLHA I FEGKITITNNRL .DCLHPSCSTSFHPNIQGAKSSSDV.KSYLDKDGDT
31 ToLCNDV   DGSPHIHV V FEGKFQCKNNRF .DLVSPSRAAHFHPNIQGAKSASDV.KAYIDKDGDV
32 AYVV      DGSPHLHV I FEGKYKCQNNRF .DLISPTRSAHFHPNIQGAKSSSDV.KSYIDKDGDT
33 TYLCSV    DGQPHLHM I FEGKFNCKNNRF .DLVSPTRSAHFHPNIQGAKSSSDV.KSYIDKDGDV
34 TYLCCNV   NGTPHLHV I FEGKFQCKNQRF .DLTSPHRSAHFHPNIQGAKSSTDV.KSYMEKDGDV
35 ToLCBaV   DGEPHLHV I FEGKFQCTNNRF .DLVSPTRSAHFHPNIQKAKSSSDV.KAYVEKDGDF
36 ToLCJV    NGEPHLHV L FEGKFQCKNNRF .DLVSPTRSAHFHPNIQGAKSSTDV.KAYVEKDGDF
37 ChiLCAV   NGEPHLHV V FEGKYQCTNNRF .DLISPTRSAHFHPNIQRAKSSSDV.KAYVEKDGDF
38 ChiLCV    NGEPHLHV V FEGKYQCTNNRF .DLISPTRSAHFHPNIQRAKSSSDV.KAYVEKDGDF
```

```
              110        120       130       140       150       160
 1 TYLCV      IDF..GVFQIDGRSARGGQQSANDAYAEALNSGNKSEALNILKEKAPKDYILQFHNL..S
 2 ToLDeV     LEW..GEFQVDGRSARGGSQTANDAAAEALNSGSKEAALQIIREKLPEKYLFQFHNL..N
 3 ToLCV-2    LEW..GEFQIDGRSARGGQQTANDACAEALNASSKEAAMQIIKEKLPEKFLFQYHNL..S
 4 ToMoWV     LEW..GEFQIDGRSARGGQQTANDAAAEALNAPSKESAMQIIREKLPEKYLFQYHNL..N
 5 ToLDV      IEW..GEFQVDGRSARGGQQTSNDAAAEALNASSKEEAMMIIKEKLPEKFLFQYHNL..S
 6 BGMV       IEW..GHFQVDGRSARGGQQTANDAASEALNASSKEEAMQIIKEKLPEKFLFQYHNL..S
 7 SiYNV      VEW..GTFQIDGRSARGGQQTANDVAAEALNAGTKEEAMMIIKEKMPEKFLFQYHNL..S
 8 ToLCPVV    VEW..GDFQIDGRSARGGQQTANDTYAKALNASSAEEALQIIKEEQPQHFFLQHHNL..V
 9 ToGLSV     CES..GRYKISG....GTKTNKDDVYHNAINATSASEALAIIRTGDPKAFIVQHHNI..S
10 SLCuV      CES..GQYKVSG....GSKSNKDDVYHNAVNAGSAGEALDIIKAGDPKTFIVNYHNL..L
11 CabLCV     CES..GQYKVSG....GTKANKDDVYHNAVNAGCVEEALAIIRAGDPKTFIVSYHNV..R
12 ToCmMV     CES..GQYKVSG....GTKANKDDVYHNAVNASSPGEALDIIRAGDPKTFLVNYHNV..K
13 ToSLCV     CES..GQYKVSG....GTKSNKDDVYHNAVNAASAGEALDIIRAGDPKTFIVSYHNV..K
14 TYMLCV     IQW..GEFQIDGRSARGGQQSANDTYAKALNARSSEEALQIIKEEQPQHFFLQHHNL..L
15 BoGMV      IDW..GEFQIDGRSARGGKQSTNDTYAKALNAASADEALQIIKEEQPQHFFLQHHNI..V
16 CdTAV      IDH..GVFQIDGRSARGGCQSANDSYAKALNAGDASQALNILREEQPRDFFANYHNL..K
17 JMV        VEW..GEFQIDGRSARGGCQSANDTYAKALNASSADEALQIIKEEQPQHFFLHHHNL..V
18 AbMV       AEW..GEFQIDGRSARGGQQTANDSYAKALNAGDVQSALNILKEEQPKDYVLQNHNI..R
19 ToVCLDeV   IDF..GVFQIDGRSARGGQQSANDTYAKVLNAGSILEALNILKEEQPKDFVLQHHNI..R
20 PSLDV      IDH..GIFQIDGRSARGGQQSANDTYAKVLNSGSVMEALNILREEQPKDFVLQHHNI..R
21 ToMoLCV    IDH..GVFQIDGRSARGGQQSANDTYAKVLNAGSVMEALNILREEQPKDFLLQHHNI..R
22 ToGLDV     CSWYFSRSTADQLEEVSNLPTTRMPRSSTLAPSQRPSIYSGRNNQKTSSFKIITYVLIYN
23 SPLCSiV-2  LTW..GEFQIDGRSARGGQQTANEAYAQALNAGNKAEALQIIKELCPKDFVLQFHNI..N
24 SPLCHnV    VDS..GVFQVDGRSARGGQQTANDAYAQALNTGCKSEALQIIKELCPKDFVLQFHNI..N
25 SPMV       LTW..GEFQIDGRSARGGQQTANDAAAEALNSGSKEAALQIIREKLPEKFIFQYHNL..C
26 SPLCLaV    IEW..GEFQVDGRSARGGQQTANDAAAEALNSGSKEAALQIIREKLPEKFIFQYHNL..C
27 MYMV       LDH..GSFQVDGRSARGGKQSANDAYAEALNSGSKLQALNILREKAPKDYILQFHNL..N
28 EACMV      LDD..GIFQVDARSARGEGQHLAQVYADALNASSKSEALQIIKEKDPKSFFLQFHNI..S
29 CLCuGV     LEW..GEFQIDGRSARGGQQTANDAYAAALNAGSKAEALRVIRELAPKDFVLQFHNL..N
30 ACMV       VEW..GQFQIDGRSARGGQQSANDAYAKALNSGSKSEALNVIRELVPKDFVLQFHNL..N
31 ToLCNDV    LEW..GVFQIDGRSARGGQQTANDAYAQAINTGNKTDALKVLKELAPKDYVLQFHNL..I
32 AYVV       LEW..GEFQIDGRSARGGQQTANDAYAQALNSGSKSEALNVIKELAPKDYVLQFHNL..N
33 TYLCSV     LEW..GTFQIDGRSARGGQQTANDAYAKAINAGRKSEALDVIKELAPRDYILHFHNI..N
34 TYLCCNV    LDH..GVFQIDGRSARGGCQSANDAYAEAINSGSKASALNILREKAPKDFVLQFHNL..N
35 ToLCBaV    IDF..GVFQIDGRSARGGQQSANDAYAEAINSGSKAAALDILREKAPKDFVLQFHNL..N
36 ToLCJV     IDF..GVFQIDGRSARGGQQSANDAYAEAINSGSKAAALNILREKAPKDYVLQFHNL..N
37 ChiLCAV    IDF..GVFQIDGRSARGGCQSANDAYAEAINSGSKSSALNILREKAPKDYVLQFHNL..N
38 ChiLCV     IDS..GVFQIDGRSARGGCQSANDAYAEAINSGSKAAALNILREKAPKDYVLQFHNL..N

                  170        180       190       200        210
 1 TYLCV      SNLDRI.... SPPLEVYVSPFLSSSFNQV DELEEWVAENVV..SSAARPWR.....PN
 2 ToLDeV     CNLDKI.... SKAPEPWSPPFPLSSFTNV EEMQEWADDYFGR.GAAARPER.....PI
 3 ToLCV-2    SNLDRI.... SKPPEQWVPPFPLSSFTHV DEMQERANDYFGR.DAAARPER.....PI
 4 ToMoWV     SNLDRI.... NKPPEPWVPPFRLSSFTNV DEMQEWVDEYFGR.GAAARPER.....PI
 5 ToLDV      SNLDRI.... KKAPDPWVPPFPLSSFTNV DEMQEWADEYFGR.GAAARPVR.....PM
 6 BGMV       SNLDRI.... TKAPDPWSPPYHLSSFTNV REMQEWADDYFGR.GAAARPER.....PI
 7 SiYNV      SNLDRI.... KKAPEPWTPPFPLSSFTNV DEMQSWADDYFGLVVSAARPER.....PV
 8 ToLCPVV    ANAQRI.... QKAPEPWAPPFPLSSFTNV EEMQEWADDYLGR.DAAARPER.....PI
 9 ToGLSV     ANIHKI.... AQSPEPWTPPFQLSSFTNV DEMQEWADDYFGR.LSAARPHFHNSLRPV
10 SLCuV      ANVERL.... QKPPEPWVPPFELSSFTSV EELQDWADDYFNE.CSAA..AR.....PT
11 CabLCV     ANIERL.... TKAPEPWAPPFQLSSFTNV DEMSSWADDYFGR.SAAARAER.....PI
12 ToCmMV     ANLERI.... QKAPAPWVPPFPLSSFTNV DEMQEWADDYFGR.DAAARPIR.....PI
13 ToSLCV     ANIERL.... QKAPEPWVPPFPLSSFTNV DEMQEWADDYFGR.DAAARPER.....PI
14 TYMLCV     ANATKI.... RKPPEPWVPPFRLSSFINV VDMQEWADDYFGR.DSAARPER.....PI
15 BoGMV      ANAARI.... RKPVSQWVPPFPISSFVNV HQMRQWADDYFGR.DAAARPER.....PK
16 CdTAV      ANAERI.... AKAPEPWIPPFPLSSFTQV EEMQEWADEFFGR.GAAARPER.....PI
17 JMV        ANAQRI.... QKAPEPWVPPFQLASFTNV EEMQEWADEYFGM.DSAARPER.....PI
18 AbMV       SNLERI.... AKAPEPWVPRFPLSSFTAV EEMQEWADDYFGS.GSAARPDR.....PL
19 ToVCLDeV   SNLERI.... QKPPEPWAPPFPLSSFTNV DEMQEWADDYFGR.SSAARPER.....PI
20 PSLDV      SNLERI.... QKAPEPFVPPFPLSSFTLV EEMQEWADSYFGL.DAAARPER.....AI
21 ToMoLCV    SNLERI.... QKAPEPWVPPFPLSSFTEV VEMQEWADDYFGM.DPAARPER.....PI
22 ToGLDV     VSLQRLRSHG LRFTSPHSIPFPTTC.KSG MII.......LG..EVPLRGRRDLLVLSS
23 SPLCSiV-2  NNLDRI.... SPPVEVYTSPFLSSSFNNV DIISNWAADNIK..DAAARPDR.....PI
24 SPLCHnV    NNLDRI.... SPPPSVYSSPFSLSSFNNV DIISDWAAENVM..DAAARPDR.....PI
25 SPMV       SNLDRI.... SPPPSVYSSPFSLSSFNNV DIISDWAAENVM..DSAARPDR.....PI
26 SPLCLaV    GNLDRI.... SPPPSVYSSPFSSSSFNAV DIISDWAAENVM..DSAARPDR.....PI
27 MYMV       CNLSRI.... ADEVPLYVSPYSLSTFDKV SYISSWASENVR..DSAARPDR.....PI
28 EACMV      ANADRI.... QAPPQTYVSPFLSSSFTNV EELEVWVSENVM..GSAARPWR.....PN
29 CLCuGV     SNLERI.... QEPPAPYVSPFLSSSFDQV EELEEWAAENVV..EAAARPSR.....PI
30 ACMV       SNLERI.... QEPPAPYISPFPCSSFDQV DELEEWVADNVR..ASAARPWR.....PN
31 ToLCNDV    NNLDRI.... QTRSEVYVSPFSISSFDRV ANLVDWAKCNVV..CSAARPLR.....PI
32 AYVV       ANLDRI.... APPLEVFVCPFLSSSFDQV EELEEWVSENVK..DAAARPWR.....PK
33 TYLCSV     SNLNMV.... QVPPAPYVSPFLSSSFDQV DELEHWVSENVM..DVAARPWR.....PV
34 TYLCCNV    SNLDRI.... TPPMEEYISPFSSSSFDQV EELDEWAVDNVV..SAAARPLR.....PV
35 ToLCBaV    ANLDRI.... TPSAEVYVSPFLCSSFDTV EELHHWAAENVV..DAAA.AIR.....PI
36 ToLCJV     SNLDRI.... TPPPEVYISPFSSSSFDQV DELEEWAAENVV..GAAARPLR.....PI
37 ChiLCAV    SNLDRI.... TPPPEVYVSPFLSSSFDQV AELEDWASENVM..GAAARPLR.....PM
38 ChiLCV     ANLDRI.... TPPPDVYVSPFLSSSFDQV GELEEWASVNVV..GAAARPLR.....PM
```

```
                   220            230            240            250            260           270
 1 TYLCV       SIVIEGDS T KTMWARSLGPHNYLCGHLDLSPKVYSNDAWYNVIDDVDPHYL..KHF E
 2 ToLDeV      SIIIEGCS T KTMWARALGPHNYLSGHLDFNSKVYSNGADYNVIDDVSPQYLKMKHW E
 3 ToLCV-2     SIIVEGDS T KTMWARALGPHNYLSGHLDFNSRVYSNEAKYNVIDDVIPDYLKKKHW E
 4 ToMoWV      SIIIEGDS T KTMWARALGSHNYLSGHLDFNSKVYSNEVEYNVIDDVTPQYLKMKHW E
 5 ToLDV       SIIVQGDS T KTMWARALGSHNYLSGHLDFNSRVYSNDVEYNVIDDVTPHYLKLKHW E
 6 BGMV        SIIIEGDS T KTMWARALGTHNYLSGHLDFNSKVFSNHAEYNVIDDIAPHYLKLKHW E
 7 SiYNV       SIIIEGDS T KTMWARALGAHNYLSGHLDFNSRVYSNEVDYNVIDDVPPHYLKLKHW E
 8 ToLCPVV     SIIIEGDS T KTMWARSLGSHNYLSGHLDFNSRVYSNDAEYNVIDDVTPQYLKMKHW E
 9 ToGLSV      SIIVEGDS T KTMWARALGSHNYLSGHLDFNSRVYSNDVDYNVIDDVAPHYLKMKHW E
10 SLCuV       SIIIEGGS T KTMWARSLGPHNYLSGHLDFNSRVFSNDVKYNVIDDVAPHYLKLKHW E
11 CabLCV      SIIVEGDS T KTMWARALGPHNYLSGHLDFNSKVFSNNAEYNVIDDIAPHYLKLKHW E
12 ToCmMV      SIVIEGNS T KTMWARALGPHNYLSGHLDFNSRVYSNDVLYNVIDDVSPHYLKLKHW E
13 ToSLCV      SIIIEGES T KTMWARALGPHNYLSGHLDFNSRVYSNEAEYNVIDDITPQYLKMKHW E
14 TYMLCV      SLIVEGDS T KTMWARALGSHNYLSGHMDFNSRVYSNDVEYNVIDDVNPQYLKIKHW E
15 BoGMV       SIIIEGDS T KTMWARALGKHNYLSGHLDFNARVYSNDALYNVIDDISPQYLKLKHW E
16 CdTAV       SIIIEGDS T KTMWARALGPHNYLSGHLDFNSRVYSNQVEYNVIDDVAPHYLKLKHW E
17 JMV         SVIVEGDS T KTMWARSLGPHNYLSGHLDFNPRVYSNDVEYNVIDDVAPHYLKLKHW E
18 AbMV        SLIVEGDS T KTMWARALGPHNYLSGHLDFNGRVYSNEVEYNVIDDVAPHYLKLKHW E
19 ToVCLDeV    SIIVEGDS T KTMWARALGPHNYLCGHLDLNSRVYSNNVDYNVIDDVNPQYL..KHW E
20 PSLDV       SIIIEGDS T KTMWARSLGVHNYLSGHLDFNSRVYSNDVEYNVIDDISPHYLKMKHW E
21 ToMoLCV     SLIIEGDS T KTMWARGLGPHNYLSGHLDFNPRVYSNEAEYNVIDDITPQYLKMKHW E
22 ToGLDV      RVIVERGR C HVLYGSTIICADTYISILGFTQMTWSITSLMMSLRNTYSY....STG N
23 SPLCSiV-2   SIVIEGPS I KTVWARSLGPHNYLCGHLDLNPRVYSNSAWYNVIDDVDPQYL..KHF E
24 SPLCHnV     SIVIEGPS I KTVWARSLGPHNYLCGHLDLSPKVYSNSAWYNVIDDVNPQYL..KHF E
25 SPMV        SIVIEGPS I KTVWARSLGPHNYLCGHLDLSPKVYSNSAWYDVIDDVNPQYL..KHF E
26 SPLCLaV     SIVIEGPS I KTVWARSLGPHNYLCGHLDLSPKVYSNSAWYNVIDDVNPQYL..KHF E
27 MYMV        SIVIEGDS T KTMWARALGPHNYLCGHLDLNSKIYSNDAWYNVIDDVDPHYL..KHF E
28 EACMV       SIVLEGDS T KTMWARSLGPHNYLCGHLDLSPKVYSNDAWYNVIDDVDPHYL..KHF E
29 CLCuGV      SIVIEGES T KTVWARSLGPHNYLCGHLDLSPKVFSNDAWYNVIDDVDPHYL..KHF E
30 ACMV        SIVIEGDS T KTIWARSLGPHNYLCGHLDLSPKVFNNAAWYNVIDDVDPHYL..KHF E
31 ToLCNDV     SIVIQGDS T KTMWARCLGPHNYLCGHLDLSPKVYSNDAWYNVIDDVDPHYL..KHF E
32 AYVV        SIVVEGES T KTMWARSLGPHNYLCGHLDLSPKVYSNAAWYNVIDDVDPHYL..KHF E
33 TYLCSV      SIVIEGDS T KTMWARSLGPHNYLCGHLDLSQKVYSNNAWYNVIDDVDPHYL..KHF E
34 TYLCCNV     SIVIEGDS T KTMWARSLGPHNYLCGHLDLSPKVYSNDAWFNVIDDVDPHYL..KHF E
35 ToLCBaV     SIVIEGDS T KTMWARSLGPHNYLCGHLDLSPRVYSNDAWYNVIDDVDPHYL..KHF E
36 ToLCJV      SIVIEGDS T KTVWARSLGPHNYLCGHLDLSPKVYSNDAWYNVIDDVDPHYL..KHF E
37 ChiLCAV     SIVIEGDS T KTMWARSLGPHNYLCGHLDLSPKVYSNDAWYNVIDDVDPHYL..KHF E
38 ChiLCV      SIVIEGDS T KTMWARSLGPHNYLCGHLDLSPKVYSNDASYNVIDDVDPHYL..KHF E

                     280            290            300            310            320            330
 1 TYLCV       FM AQRDWQSNTKYGKPIQI GGI TIFLCNP PTS YREYLDEEKNIS KNWALKN TF
 2 ToLDeV      LI AQKDWQSNCKYGKPIQI GGI SIVLCNP EGQ YKDFLEKEENAA RAWTLHN KF
 3 ToLCV-2     LI AQKDWQSNCKYGKPVQI GGI SIVLCNP EGS YKTFLEREENVS KNWTLHN KF
 4 ToMoWV      LI SQRDWQSNCKYGKPVQI GGI SIVLCNP EGA YKEFLDKHENSS KAWTLHN KF
 5 ToLDV       LI AQKDWQSNCKYGKPVQI GGI SIVLCNP EGA YKDFLEKEENAS KSWTLYN KF
 6 BGMV        LM AQKDWQSNCKYGKPVQI GGI SIVLCNP EGA YKCFLDKEENAA KNWTIHN KF
 7 SiYNV       LI AQKDWQSNCKYGKPVQI GGI SIVLCNP EGS YKISIMFRNQGR REKGVFW KR
 8 ToLCPVV     LI AQRDWQSNCKYGKPVQI GGI SIVLCNP EGS YKDFLEKQENVS KNWTLHN KF
 9 ToGLSV      LI AQRDWQSNCKYGKPVQI GGI SIVLCNP EGS YKEFLDKEENLS KNWTLHN KF
10 SLCuV       LI AQRDWQSNCKYGKPVQI GGI SIVLCNP EGS YKDFLDKAENSA REWTEKN KF
11 CabLCV      LI AQRDWQSNCKYGKPVQI GGI SIVLCNP EGS YISFLNKEENAS RAWTTKN KF
12 ToCmMV      LI AQIDWQSNCKYGKPVQI GGI SIVLCNP EGA YKDFLAKEENAS KSWTLHN KF
13 ToSLCV      LI AQKDWQSNCKYGKPVQI GGI SIVLCNP EGA YKSFLEKEENKA KDWTLHN KF
14 TYMLCV      LI AQKDWQSNCKYGKPVQI GGI AIVLCNP EGS YKRYLDKEENTS RAWTLHN KF
15 BoGMV       LI AQRDWQSNCKYGKPVQI GGI SIVLCNP EGA YQDFLNKWENVA KNWTEHN TF
16 CdTAV       LI SQKDWQSNCKYGKPVQI GGI SIVLCNP EGS YKDFLNKEENAS RNWTLKN LF
17 JMV         LL AQRDWQSNCKYGKPVQI GGI AIVLCNP EGS YKDFLDKEENSS RNWTIKN VF
18 AbMV        LL AQKDWQSNCKYGKPVQI GGI AIVLCNP EGS YKEYLDKEENTG RNWTLKN IF
19 ToVCLDeV    II AQRDWQSNCKYGKPVQI GGI SIVLCNP AGS YKAFLDKEENAS RAWTLHN KF
20 PSLDV       LI AQRDWQSNCKYGKPVQI GGV SIVLCNP EGS YKDFLDKEENAS RNWTLRN QF
21 ToMoLCV     LI AQKDWQSNCKYGKPVQI GGI SIVLCNP EGA YKDFLDKEENTA KNWTLHN KF
22 ToGLDV      CL PKGTGNQIASTASQFKL EAY QLCFAIQ EGS YKEFLEKQENVS NDWTIHN KF
23 SPLCSiV-2   FI AQKDWQSNTKYGKPVQI GGI TIFLCNP EGS FKSWLDKTEQDA RQWACKN VF
24 SPLCHnV     FM AQKDWQSNCKYGKPVQI GGI TIFLCNP EGS FKLWLDKPEQGA KIWATAN LF
25 SPMV        FM AQKDWQSNCKYGKPVQI GGI TIFLCNP EGS FKLWLDKPEQEA KNWAVKN IF
26 SPLCLaV     FM AQKDWQSNCKYGKPVQI GGI TIFLCNP EGS FKLWLDKPEQGA KNWATAN IF
27 MYMV        FM AQRDWQSNVKYGKPTHI GGI TIFLCNP PKS YKEYLDEADNTA KLWASKN EF
28 EACMV       FM AQRDWQSNTKYGKPIQI GGI TIFLCNP PTS YKEFLDEEKNQS KAWALKN TF
29 CLCuGV      FM AQKDWQSNTKYGKPVQI GGI TIFLCNP PNS YKEYLDEDKNAH KSWALKN TF
30 ACMV        FM SQRDWQSNTKYGKPVQI GGI TIFLCNP PTS YKEFLDEEKQEA KAWALKN IF
31 ToLCNDV     FM AQRDWQSNTKYGKPVMI GGI TIFLCNE PNS YKEYLDEEKNAA KQWAIKN VF
32 AYVV        FM AQRDWQSNTKYGRPIQI GGI TIFLCNP PTS YKEYLEEEKNSA KAWAIKN EF
33 TYLCSV      FM SQRDWQSNTKYGKPIQI GGI TIFLCNP PQS FKEYLDEEKNQT KNWAIKN IF
34 TYLCCNV     FM AQRDWQSNTKYGKPVQI GGI TIFLCNP PNS YKEYLEEEKNSA RNWAVKN IF
35 ToLCBaV     FM AQRDWQSNTKYGKPVQI GGI TIFLCNP PNS YKEFLDEDKNNA KQWALKN TF
36 ToLCJV      FM AQRDWQSNTKYGKPVQI GGI AIFLCNP PNS YKEFLDEEKNSA KNWALKN TF
37 ChiLCAV     FM AQRDWQSNTKYGKPVQI GGI AIFLCNP PNS YKEFLDEEKNSA KHWALKN TF
38 ChiLCV      FM AQRDWQSNTKYGKPVQI GGI AIFLCNP PNS YKEFLDEEKNSA KHWALKN TF
```

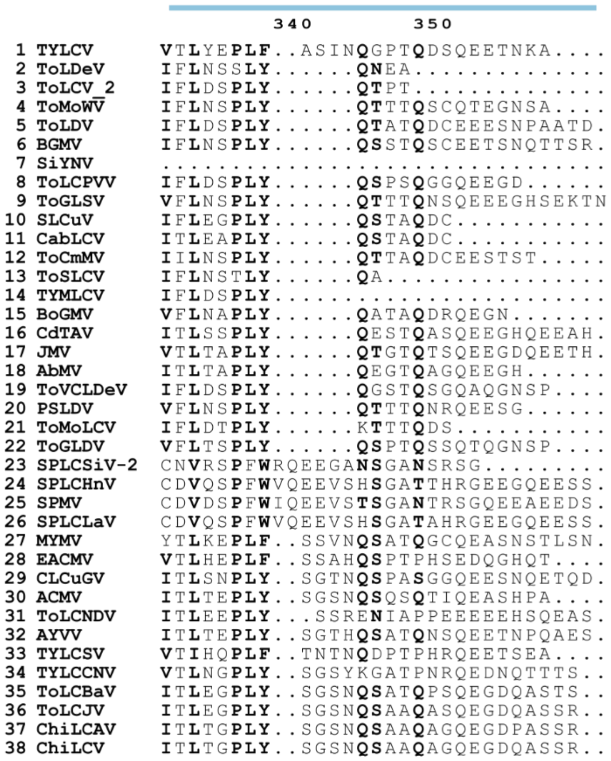


**Figure S5. Rep proteins from selected begomoviruses**. Multiple sequence alignment of the Rep proteins from selected begomoviruses from Figure 3A using MUSCLE. Alignment was represented with Espript 3.2 (Robert *et al*, 2025).

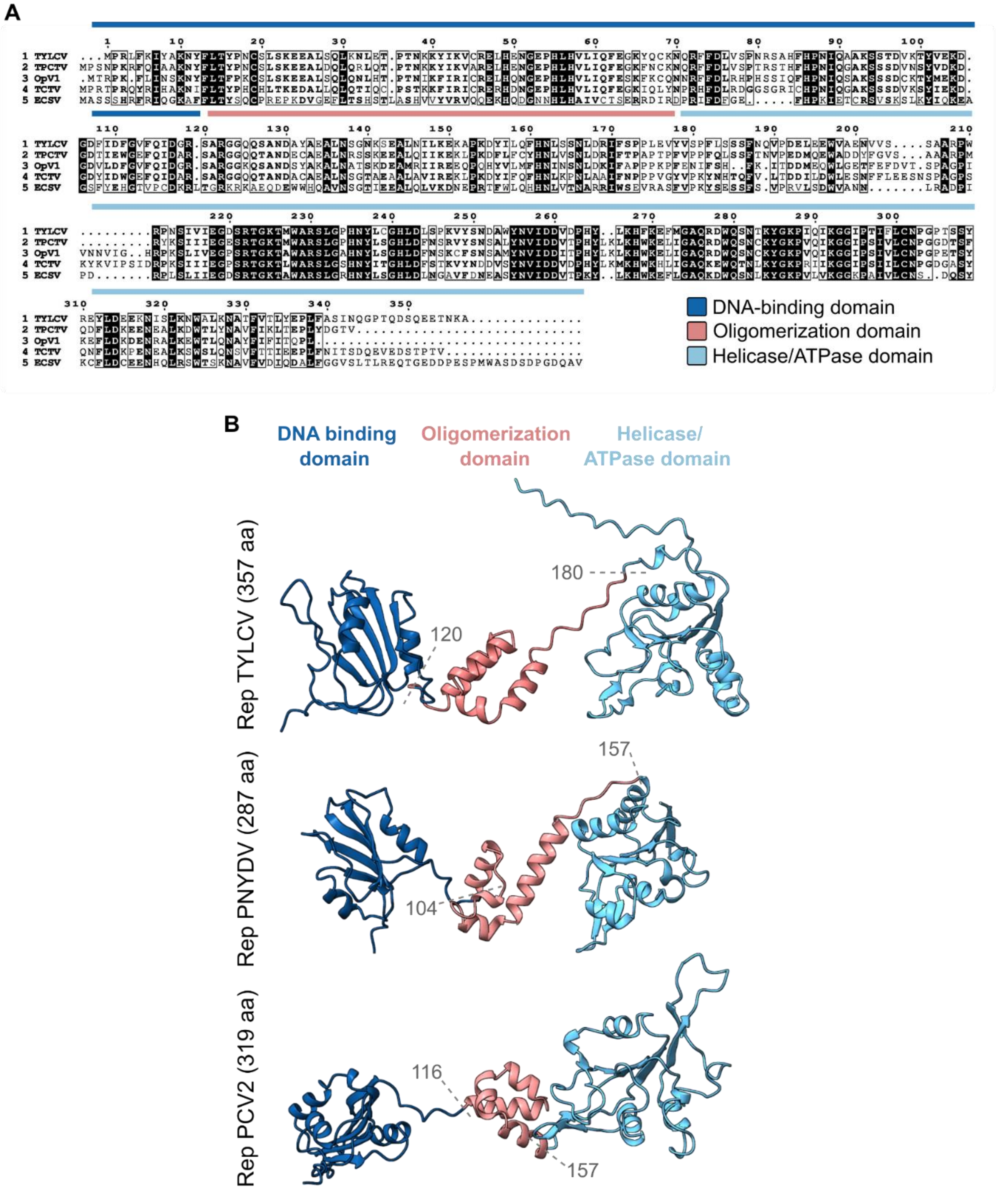


**Figure S6. Rep proteins from selected geminiviruses and related CRESS viruses**. (A) Multiple sequence alignment of the Rep proteins from selected geminiviruses from Figure 4A,B using MUSCLE. Alignment was represented with Espript 3.2 (Robert *et al*, 2025).

(B) Structure and domain organization of the Rep proteins from PNYDV and PCV2. For PNYDV, prediction was performed using Alphafold3 (https://alphafoldserver.com/). PCV2 structure and domain organization are described in Tarasova *et al* (2021). A comparison with

Rep from TYLCV, as well as the position (in aa) of the different domains are shown. Rep conformation for PNYDV and PCV2 is shown after alignment with the Rep from TYLCV in ChimeraX.
